# GABA(A) receptor activation drives GABARAP-Nix mediated autophagy to radiation-sensitize primary and brain-metastatic lung adenocarcinoma tumors

**DOI:** 10.1101/2023.11.29.569295

**Authors:** Debanjan Bhattacharya, Riccardo Barille, Donatien Kamdem Toukam, Vaibhavkumar S. Gawali, Laura Kallay, Taukir Ahmed, Hawley Brown, Sepideh Rezvanian, Aniruddha Karve, Pankaj B. Desai, Mario Medvedovic, Kyle Wang, Dan Ionascu, Nusrat Harun, Chenran Wang, Andrew M. Baschnagel, Joshua A. Kritzer, James M. Cook, Daniel A. Pomeranz Krummel, Soma Sengupta

## Abstract

In non-small cell lung cancer (NSCLC) treatment, targeted therapies benefit only a subset of NSCLC, while radiotherapy responses are not durable and toxicity limits therapy. We find that a GABA(A) receptor activator, AM-101, impairs viability and clonogenicity of NSCLC primary and brain metastatic cells. Employing an *ex vivo* ‘chip’, AM-101 is as efficacious as the chemotherapeutic docetaxel, which is used with radiotherapy for advanced-stage NSCLC. *In vivo*, AM-101 potentiates radiation, including conferring a survival benefit to mice bearing NSCLC intracranial tumors. GABA(A) receptor activation stimulates a selective-autophagic response via multimerization of GABA(A) Receptor-Associated Protein (GABARAP), stabilization of mitochondrial receptor Nix, and utilization of ubiquitin-binding protein p62. A targeted-peptide disrupting Nix binding to GABARAP inhibits AM-101 cytotoxicity. This supports a model of GABA(A) receptor activation driving a GABARAP-Nix multimerization axis triggering autophagy. In patients receiving radiotherapy, GABA(A) receptor activation may improve tumor control while allowing radiation dose de-intensification to reduce toxicity.

**Highlights:**

- Activating GABA(A) receptors intrinsic to lung primary and metastatic brain cancer cells triggers a cytotoxic response.
- GABA(A) receptor activation works as well as chemotherapeutic docetaxel in impairing lung cancer viability *ex vivo*.
- GABA(A) receptor activation increases survival of mice bearing lung metastatic brain tumors.
- A selective-autophagic response is stimulated by GABA(A) receptor activation that includes multimerization of GABARAP and Nix.
- Employing a new nanomolar affinity peptide that abrogates autophagosome formation inhibits cytotoxicity elicited by GABA(A) receptor activation.

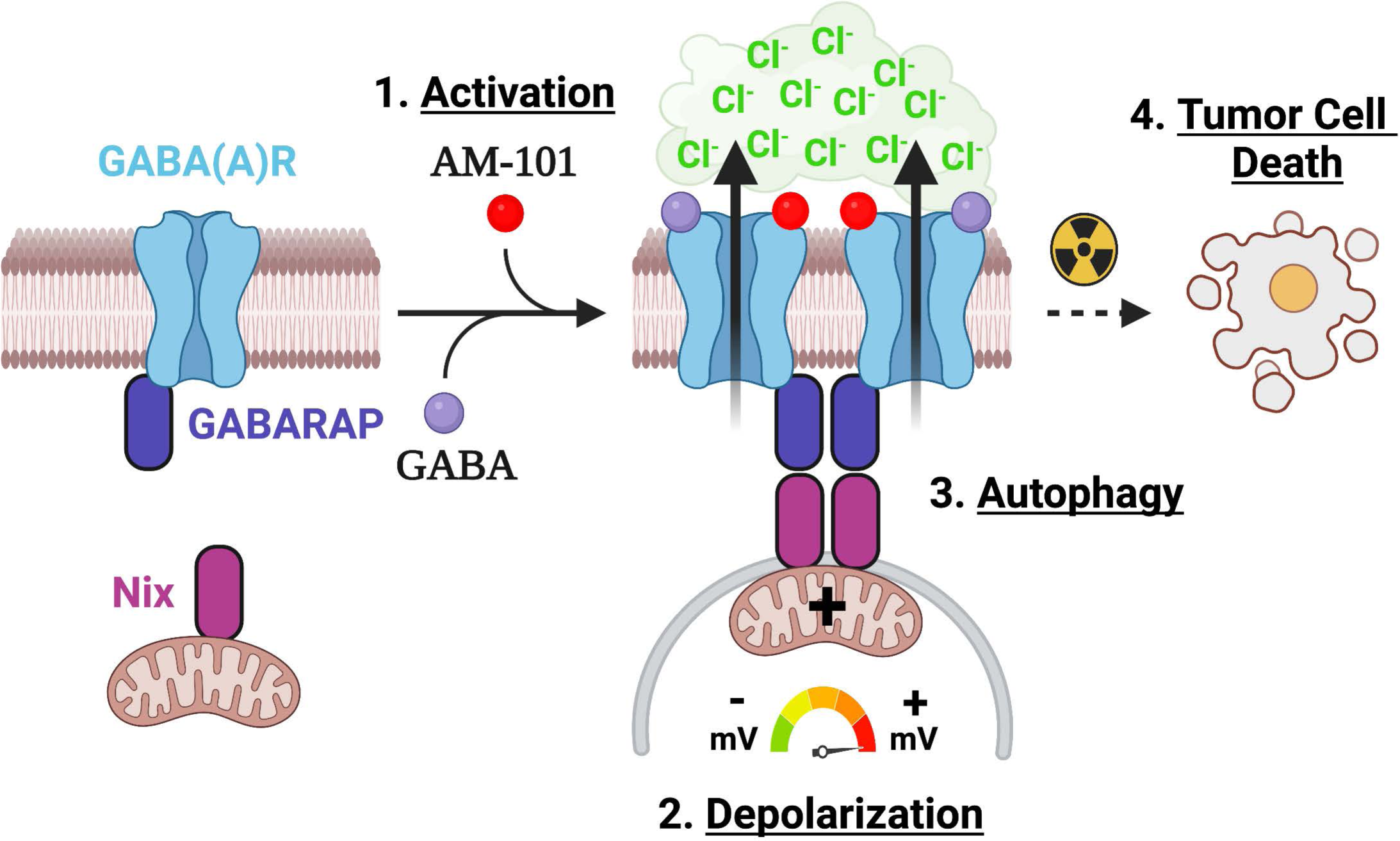

## Introduction

Lung cancer is the leading cause of cancer associated mortality in the United States, with approximately 350 deaths per day.^1^ The most common lung cancer type, non-small cell lung cancer (NSCLC), comprises 80-85% of lung cancers and has an overall 5-year survival rate of 64%, but that drops to 37% once tumor cells invade the lymphatic system.^1^ Most NSCLC patients will clinically present with metastatic disease, which is usually incurable.

A highly challenging metastatic site to treat is the brain, which accounts for 30% of NSCLC metastatic cases.^2^ However, post-mortem analysis of NSCLC patients suggests incidence of brain metastasis is significantly higher.^3^ Standard-of-care for lung cancer brain metastasis includes surgical resection for large solitary lesions and stereotactic brain radiosurgery (SRS) in most other cases involving less than ten lesions. Otherwise, whole brain radiation therapy (WBRT) is employed. However, both SRS and WBRT are associated with toxicity including radiation necrosis and neurocognitive effects. Managing potential toxicities and overcoming ensuing radiation resistance pose major challenges. While cranial radiotherapy is usually part of a multimodal treatment regimen for NSCLC brain metastasis, the median survival is only about 7-10 months.^4–6^ To enhance radiation effectiveness the plant alkaloid docetaxel (Taxotere) is commonly employed as a chemotherapeutic in conjunction with radiotherapy for advanced-stage NSCLC.^7^ Docetaxel, however, is associated with significant co-morbidities, including weakened immunity and peripheral neuropathy. Furthermore, docetaxel is not employed for treatment of brain metastasis, as it is not brain-penetrant.

Significant attention has recently been focused on targeting as an anti-cancer stratagem autophagy, the phenomenon of self- or auto-degradation. Both agents that inhibit or induce autophagy are being explored to impair cancer cell viability. Agents that inhibit autophagy by interfering with the assembly of the double-layered autophagosomes include the immunosuppressive anti-parasitic drugs chloroquine and hydroxychloroquine.^8^ Recently, cell-penetrant peptides have been engineered that inhibit autophagy more selectively by blocking autophagy-specific protein-protein interactions.^9^ Agents which appear to induce autophagy include the anti-diabetic metformin and mTor inhibitors (everolimus, temsirolimus, and rapamycin). In support of an autophagic induction anti-cancer approach, recent reports suggest that enhanced expression of key autophagy associated proteins prolong patient survival.^10,11,12^ Induction of autophagy by the plant-derived compound methyl jasmonate has been found to impart therapeutic effects in cultures of human NSCLC cells.^13^ As well as molecules inducing autophagy, radiation is reported to enhance expression of autophagy associated proteins.^14–16^

While investigation of how stress induces autophagy is under intense study, leveraging the stress response vis a vis autophagy as an anti-cancer approach remains poorly explored. For example, induction of autophagy is often elicited in response to mitochondrial stress.^17,18^ Mediating the stress response are membrane associated proteins that signal to alter a cellular metabolic state. Prominent membrane-associated proteins sensing environmental cues are the ligand-gated ion channels, which represent a significant pharmacologic target for remediation of neurological disorders but remain poorly explored as therapeutic vulnerabilities in cancer. Amongst ion channels, the Type-A GABA receptors are a principal target. GABA(A) receptors function as the major inhibitory neurotransmitter receptors and hyperpolarize post-synaptic mature neurons in response to the binding of its agonist, the metabolite GABA. We and others have reported on enhanced expression of GABA(A) receptor subunits in disparate cancers^19,20^ and shown that GABA(A) receptors intrinsic to cancer cells are functional i.e., GABA responsive. Importantly, GABA(A) receptors in cancer cells are depolarizing,^21^ which reflects an efflux of chloride anions like the embryonic receptor. The presence of GABA(A) receptors in cancer cells and its depolarizing effect when ‘activated’, in contrast to the adult receptor in non-cancer cells, prompted us to investigate the receptor as a therapeutic vulnerability to cancer cells.^20^ This led to identification of a class of brain-penetrant molecules that activate GABA(A) receptors and impair the viability of cancer cells as well as mediate tumor control in mouse models of these cancers. We also reported that a member of this class of activators, AM-101, potentiates radiation in a mouse melanoma tumor model.^22^

Study of AM-101’s mode-of-action revealed that it elicits mitochondrial dysregulation, manifest as a depolarization of the mitochondrial transmembrane potential of cancer cells as well as activation of the intrinsic (mitochondrial) apoptotic pathway.^21^ Mitochondrial dysregulation induces lysosomal-dependent autophagy selectively in cancer cells^23,24^ and thus, suggests a potential role for autophagy in AM-101’s mode-of-action. In support of such a role, the protein GABARAP contributes to both GABA(A) receptor function and autophagy. GABARAP is reported to mediate GABA(A) receptor subunit transport to the extracellular membrane, where it aids assembly as well as multimerization of the receptor.^25–27^ The multimerization of GABA(A) receptors is proposed to create a strong localized distribution of charge and enhanced activity, as observed experimentally.^26^ Interestingly, it is theorized that binding of GABRAP to GABA(A) receptors would also enhance chloride anion conductance.^28^ While in autophagy, GABARAP is reported to multimerize and colocalizes with the autophagosome-associated protein LC3-II, serving to nucleate autophagosome assembly as well as lysosomal fusion of autophagosome and cargo degradation.^26,27,29,30^ GABARAP dimers also complex with dimers of the outer mitochondrial membrane protein BNIP3L/NIX or Nix, which is a crucial molecule in the selective-autophagic response for removal of stressed or damaged mitochondria.^29^ These findings suggest that GABARAP may be a key intermediary molecule in regulating autophagy in response to ionic perturbation as mediated by GABA(A) receptors.

Herein, we have evaluated GABA(A) receptor activation as an approach in conjunction with radiation to treat both primary and metastatic brain NSCLC. We find that GABA(A) receptor activator AM-101 is as effective as docetaxel. In both heterotopic and intracranial NSCLC mouse models, the administration of AM-101 plus radiation leads to tumor control and in the case of mice bearing an intracranial tumor, prolonged survival. Mechanistically, AM-101 acts to drive a selective-autophagic response that is dependent on GABARAP-Nix complex formation. In addition, key proteins to assembly of autophagosomes show enhanced expression when radiation and AM-101 are combined. Activating GABA(A) receptors represent a new paradigm to therapeutically treat lung cancer, most significantly brain metastatic. We present a new class of small molecule brain-penetrant autophagy inducers that conceptually support this strategy.

## Results

### Lung adenocarcinomas express GABA(A) receptor subunits

GABA(A) receptors are ligand-gated anion channels which move chloride anions across the extracellular membrane in response to the binding of its natural agonist and metabolite, GABA (Figure 1A). GABA(A) receptors form pentameric assemblies from a combination of the translation product(s) of nineteen possible *GABR* genes.^31^ The canonical hetero-pentameric receptor has an α_2_-β_2_-γ stoichiometry with subunits primarily sampled from six α (*GABRA1-6*), three β (*GABRB1-3*), and three γ’s (*GABRG1-3*) (Figure 1A). Analysis of The Cancer Genome Atlas Program (TCGA) cohort of primary lung cancer patient tumors across NSCLC subtypes squamous cell carcinoma and adenocarcinoma shows expression of *GABR* genes. All squamous cell carcinoma tumors express a distinct set of *GABR* genes, not expressed in normal tissue, including *GABRE* and *GABRA3* (Figure S1A); while adenocarcinomas show a distinct expression signature for a subset of patients, otherwise expression of *GABR* genes is broader (Figure S1B). Western blots of human lung adenocarcinoma primary tumor cell lines (H1792, A549, H1703, H460) and patient-derived brain metastatic lung adenocarcinoma cell lines (UW-lung-2 and UW-lung-16) reveals that all express *GABRA5* (Figure 1B), concordant with RT-PCR analysis (Figure S1C). RT-PCR analysis also revealed variable levels of expression of GABRA3 mRNA in all NSCLC cell lines (Figure S1D). Furthermore, immunohistochemistry (IHC) of sections from both human lung adenocarcinoma primary and matching secondary (brain metastatic) tissue shows expression of *GABRA5* (Figure 1C) and *GABRA3* (Figure S1E).

**Figure 1.**
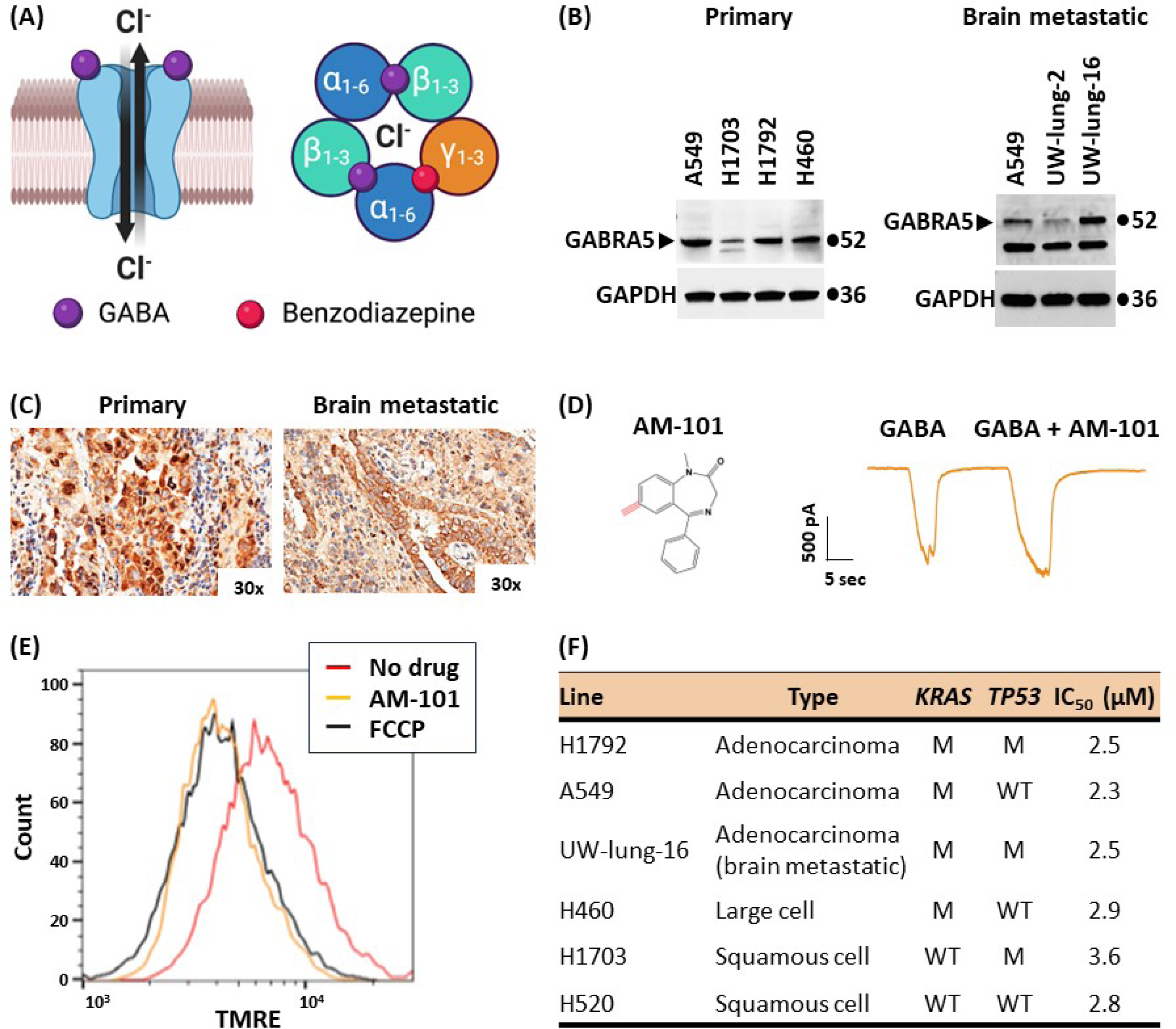
Activation of GABA(A) receptors triggers cell depolarization and death. (A) Type-A GABA (GABA(A)) receptors are ligand-gated chloride anion channels. Left, GABA(A) receptors move chloride anions (Cl^−^) out of the cell during embryonic stages of development but into the cell in mature or developed stages and are thereby depolarizing or hyperpolarizing, respectively. Right, GABA(A) receptors form hetero-pentameric structures with an α2β2γ1 stoichiometry. Two molecules of GABA (purple spheres) bind at the α-β interfaces to ‘activate’ receptor function. One molecule of benzodiazepine (red sphere) binds at the α-γ interface to enhance flow of chloride anions. (B) Left, GABRA5 or α5 protein, as shown by Western blotting of an SDS (4-15% gradient) gel, is present in NSCLC patient-derived primary cell lines representing three histological subtypes (adenocarcinoma, A549, H1792; squamous cell, H1703; large cell, H460). Right, GABRA5 or α5 protein is retained in patient-derived lung adenocarcinoma brain metastatic cell lines (UW-lung-2 and −16), as shown by Western blotting of an SDS (4-15% gradient) gel. GAPDH is used as a loading control. (C) NSCLC primary (left) and brain metastatic (right) patient tumor tissue from the same patient (or matched) stains for GABRA5 or α5 protein, as shown by immunohistochemistry staining at 30x magnification. (D) AM-101 (274.32 g/mol) is a benzodiazepine analog (left). Shown is a representative single cell patch clamp electrophysiology trace of patient-derived adenocarcinoma lung cell line H1793 (right). Cells are responsive to GABA or electro-physiologically functional. Perfusion of cells with AM-101 plus GABA elicits an enhanced response, indicating that GABA(A) receptors are benzodiazepine-responsive or ‘activated’. Representative raw current trace recording with the following parameters: GABA, 1 μM; AM-101, 4 μM. (E) Lung adenocarcinoma (H1792) cells incubated with AM-101 are depolarized, as assessed by the TMRE assay and Fluorescence Activated Cell Sorting (FACS) analysis. Shown is the degree of depolarization relative to DMSO alone treatment and FCCP, which provide negative and positive controls in this experiment, respectively. Parameters: AM-101, 2 μM; FCCP, 10 μM. (F) Half-maximal inhibitory concentration (IC50) values of patient-derived lung primary and brain metastatic lines representative of three histological lung cancer subtypes, as measured using a viability (MTS) assay and AM-101. Indicated is the *KRAS* and *TP53* mutational status of lines, where: M, mutant; WT, wild-type.

### Activation of GABA(A) receptors is depolarizing and triggers cancer cell death

Although enhanced gene expression and increased protein abundance of GABA(A) receptor subunits suggests presence of a functional receptor, it is not confirmative. We employed patch clamp electrophysiology of single primary patient-derived adenocarcinoma cells to demonstrate intrinsic, functional GABA(A) receptors. We observe a current signal in response to GABA (Figure 1D). Treatment with benzodiazepine sensitizes GABA(A) receptors in triggering a functional response leading to a potentiation effect of GABA. We observe an enhanced response to GABA plus the benzodiazepine analog AM-101 (1 and 4 μM, respectively) over GABA (1 μM) alone in H1792 cells (Figure 1D). This functional analysis reveals that H1792 cells possess: (1) intrinsic functional GABA(A) receptors; and (2) receptors that form a canonical assembly with an α-γ interface, given that a benzodiazepine can bind and elicit a positive response.

The benzodiazepine analog employed in our studies, AM-101, is an experimental drug. We have previously reported anti-tumor effects of this compound in multiple cancer settings including medulloblastoma and melanoma.^21,32^ In a previous study we reported that GABA(A) receptor activation by AM-101 leads to an efflux of chloride anions across the extracellular membrane, which contributes to a depolarization of the mitochondrial transmembrane.^21^ Similarly, we tested if AM-101 creates a shift in electric charge distribution of a lung adenocarcinoma cell type. To do so we employed the cationic fluorescent dye Tetramethyl rhodamine, ethyl ester (TMRE) and monitored its binding by Fluorescence-Activated Cell Sorting (FACS).^33^ Firstly, compared to the DMSO treated (control) cells, AM-101 causes a shift in electric charge distribution (depolarization) of H1792 cell mitochondria. Secondly, this effect occurs rapidly, within 15 min of AM-101 treatment (Figure 1E). Significantly, AM-101 induced depolarization is to an extent equivalent to that induced by treating H1792 cells with the potent mitochondrial oxidative phosphorylation uncoupler FCCP (2-[2-[4-(trifluoromethoxy)phenylhydrazinylidene]-propanedinitrile). AM-101 induces a rapid and significant depolarization of lung adenocarcinoma cells.

Membrane depolarization triggers cell death via activation of the intrinsic (mitochondrial) apoptotic pathway.^34^ Indeed, we have reported this phenomenon in cell lines of the pediatric brain cancer medulloblastoma and melanoma.^21,22^ We tested if AM-101 similarly impaired the viability of lung adenocarcinoma cells. AM-101 impairs the viability of not only lung adenocarcinoma cells, including *KRAS*- and *TP53*-mutated, but of patient-derived cells of other NSCLC subtypes, including squamous cell and large cell, with IC_50_ values in the range of 2 - 4 μM (Figures 1F and S2A). Cell proliferation impairment occurs after a 72 hr incubation with AM-101 which correlates with key events in autophagy, as will be detailed below. We also determined the IC_50_ of AM-101 for the patient-derived lung adenocarcinoma brain metastatic cells UW-lung-16^35^ and report a value like that observed for adenocarcinoma primary cells (Figures 1F, S2A, and S2B). Importantly, a single 7-ethinyl in the 1,4-benzodiazepine ring system of AM-101 bestows this cytotoxic response (Figure 1D and S2C), as diazepam (Valium) is not cytotoxic to lung adenocarcinoma cells, even at exceedingly higher concentration (Figures S2C and S2D).

### GABA(A) receptor activation potentiates radiation

Since AM-101 is cytotoxic to lung cancer cells, albeit in the low micromolar range (2 - 3 μM), we further investigated its potency and whether it was as efficacious as the anti-microtubular agent docetaxel (Taxotere), a chemotherapeutic commonly employed clinically as a radiation-sensitizer for advanced-stage NSCLC. Firstly, we employed an *ex vivo* micro-scale ‘chip’ composed of human-derived alveolar lung, endothelial lung, and GFP-tagged H1792 cells to provide an assay with a readout of efficacy that to a degree recapitulates the lung tumor microenvironment (Figure 2A). ‘Chips’ co-cultured with H1792 cells exhibit staining for phalloidin (epithelial cell marker) and PECAM1 (endothelial cell marker) (Figures S3A and S3B). H1792-GFP cells grow and spread through the epithelial compartment of the ‘Chip’ over time (Figures S3C and S3D). H1792-GFP cells die in a docetaxel concentration-dependent manner, as measured by a decrease in the number of GFP positive cells (Figures S3E and S3F). This analysis revealed that AM-101 is equally as efficacious as docetaxel in terms of its effect on H1792 cell killing and more effective at a significantly lower concentration (Figure 2B).

**Figure 2.**
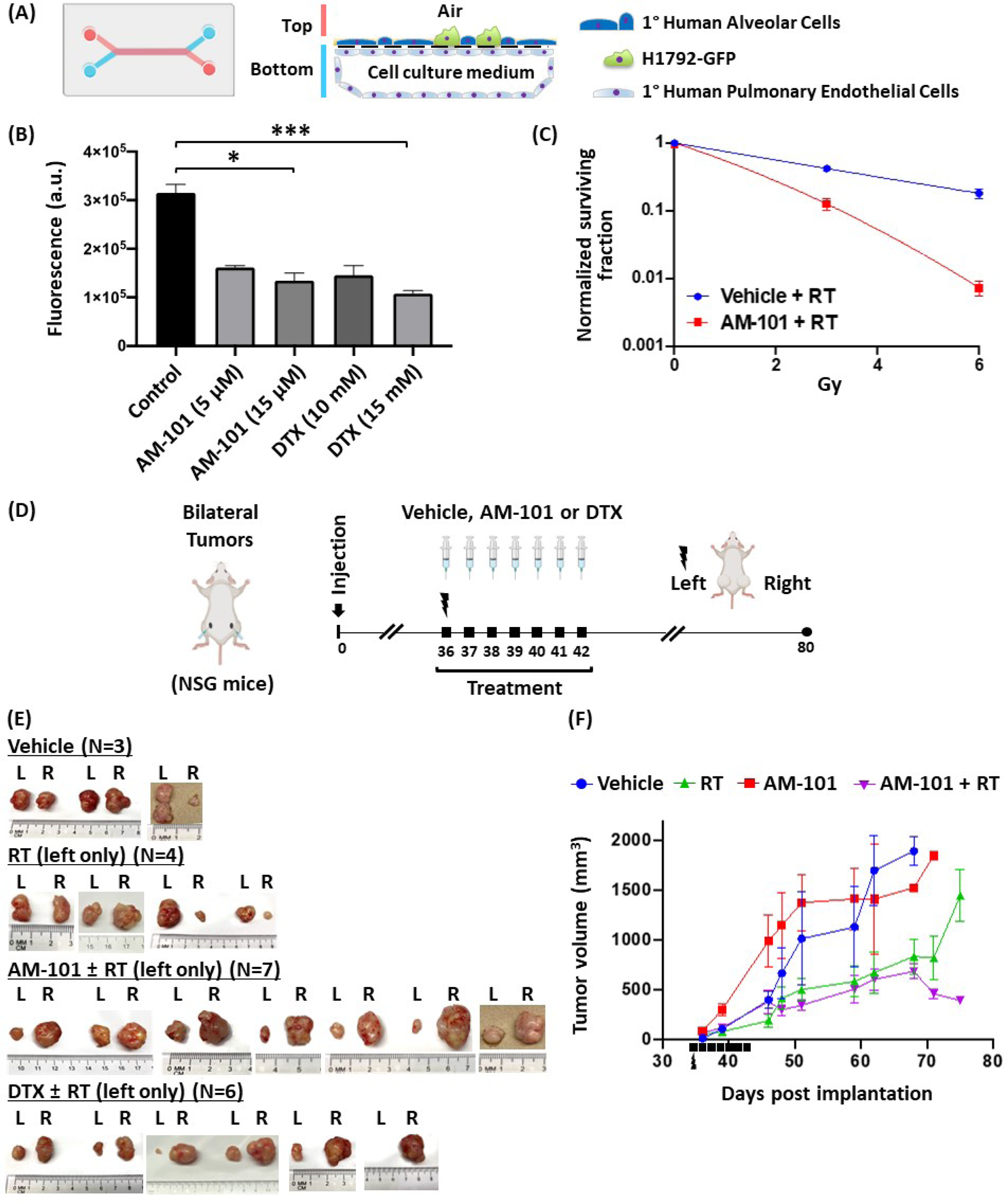
GABA(A) receptor activation potentiates radiation. (A) Illustration of a human-relevant *ex vivo* ‘chip’ employed to test AM-101 and docetaxel efficacy. Lung adenocarcinoma cancer cells (H1792-GFP, green) can be co-cultured with primary human alveolar and pulmonary endothelial cells and exposed to air (air-liquid interface) on-Chip. Cancer cells form clusters that grow and spread through the epithelial compartment of the Chip over time (Figure S3). (B) Testing of AM-101 and docetaxel *ex vivo* or ‘on-chip’ reveals that AM-101 is as cytotoxic as docetaxel, but at a significantly lower concentration. (C) A clonogenic assay was employed to examine AM-101 cytotoxicity when combined with radiation versus radiation alone. AM-101 (2.5 μM) plus radiation (RT, 3 Gy) elicits a more significant response than RT alone on colony formation in primary lung adenocarcinoma (H1792) cells. Data is presented as mean of number of colonies formed after each treatment. Vehicle or Control is DMSO, which is the vehicle of AM-101. (D) Schematic of the efficacy experiment in H1792 subcutaneous heterotopic bilateral xenograft tumors generated in NSG mice. Mice in vehicle or drug treatment groups received i.p.: vehicle, AM-101 (2.5 mg/kg), or docetaxel (DTX) (8 mg/kg) on day 36 post-implantation and then six injections once per day. Mice in RT or combo groups, received a single fraction of radiation (5 Gy) to left flank only at 2 hrs before vehicle or drug on the first day of treatment. (E) At experimental endpoint, tumors from left and right flank were resected. Shown are subcutaneous H1792 xenograft tumors in mice from different treatment groups: vehicle, RT, AM-101 ± RT, DTX ± RT. (F) Tumor volume of left and right flank tumors was measured over time. Shown is a graphical presentation of mice tumor volume when treated with vehicle, RT, AM-101, and AM-101 plus RT. Points on curve are mean tumor volume after treatment. SE. P < 0.001.

We then explored the ability of AM-101 to function as a radiation-sensitizer of lung adenocarcinoma cells. AM-101 was found to rapidly permeate the blood-brain barrier in 30 minutes without side-effects, including neuropathic pain, in rodents and monkeys.^36^ Using a clonogenic assay with lung adenocarcinoma H1792 cells, we observe a significant effect of AM-101 alone on clonogenicity (Figure S4A). But AM-101 plus radiation has a greater inhibitory effect on H1792 cell colony formation than either radiation or AM-101 alone (Figures 2C and S4A).

This greater potency of AM-101 plus radiation provided the impetus to pursue mouse efficacy studies of AM-101 alone and in combination with radiation. We generated heterotopic xenograft bilateral (left and right flank) tumors in NOD SCID Gamma (NSG) mice using H1792 cells (Figure 2D), as this adenocarcinoma cell line produces tumors that are poorly responsive to radiation.^37^ Our treatment protocol involved administering a single i.p. dose of AM-101 (2.5 mg/kg) or docetaxel (8 mg/kg) for seven consecutive days, once the tumor was palpable. For those mice in radiation receiving treatment groups, a single dose of radiation (5 Gy) was administered on the left flank only on the first day of AM-101 or docetaxel i.p. injection.

We analyzed tumor weight at experimental endpoint as well as tumor growth delay over time by measuring the tumor volume. Gross visual inspection of left (irradiated) vs right (non-irradiated) resected tumors highlights: (1) radiation alone does not exhibit a clear consistent effect on final tumor size; (2) AM-101 or docetaxel alone (right flanks) do not contribute to significant tumor control relative to vehicle; (3) combined treatment of AM-101 or docetaxel with radiation (left flanks) lowered the mean weight of the xenograft tumor in mice at experimental endpoint and resulted in consistently smaller tumor sizes (Figures 2E and S5A). In addition, there is no significant difference between median days to endpoint of control-vehicle group mice versus those receiving radiation while there is a significant difference for mice in the combined (AM-101 plus radiation group) (Figure S5B). Docetaxel also results in a statistically similar effect when combined with radiation (Figure S5A). Thus, both therapeutic agents appear to radiation-sensitize equally as well. Indeed, this is the basis for why docetaxel has been employed clinically as a radiation-sensitizer. The analysis of tumor growth delay over time reveals that AM-101 plus radiation delayed mice tumor growth compared with vehicle control, drug alone, or radiation alone (Figure 2F). This occurs even though treatment involves only a single radiation dose with seven days of AM-101 administration early in the appearance of tumor formation. The slope of the curves of the radiation versus combined treatment groups appear similar up to ∼ 68 days, at which point they diverge, suggesting that combined treatment has a protracted impact on tumor growth. This effect is manifest as an increase in median days to endpoint for those mice receiving the combination treatment, AM-101 plus radiation (Figure S5B).

### Increased survival of mice bearing a lung brain metastatic tumor

The analysis of AM-101 and docetaxel in the context of heterotopic tumors revealed that each works equally as well in potentiating radiation. As highlighted in the introduction, there is a significant demand for radiation-sensitizing agents that are brain-penetrant and non-toxic to improve standard-of-care for lung brain metastasis treatment.^6^ Docetaxel is not brain-penetrant and therefore not employed to treat lung brain metastasis. Docetaxel is also associated with significant co-morbidities including neuropathic pain.

As discussed, and shown above, GABA(A) receptors are present in both patient adenocarcinoma primary and brain metastatic lesions. We therefore explored if AM-101 is effective in potentiating radiation in the brain setting. We first examined the effect of AM-101 on clonogenicity of a patient-derived brain metastatic line, UW-lung-16. AM-101 alone impairs the viability of these cells and significantly inhibits their clonogenicity (Figure 3A). When combined with radiation, the effect is ∼ 2-fold better than AM-101 alone. Second, we generated UW-lung-16 luciferase-tagged cells from the parental cell line and employed this line for *in vivo* experiments. We generated intracranial xenograft tumors by stereotaxic injection in athymic nude mice and monitored tumor growth by bioluminescent imaging (BLI), symptoms of mice, and overall survival (Figure 3B). This experiment entailed a treatment consisting of seven-day consecutive i.p. injections of AM-101 (5 mg/kg/day) and a five-day consecutive whole-brain radiation exposure (2.5 Gy/day), overlapping with the first five days of AM-101 administration (Figure 3B). In control mice treated with vehicle (a co-solvent formulation) tumors develop within 18 days post-implantation, as visualized by BLI (Figure 3C). Tumors caused severe neurologic symptoms in mice by day 26, which manifest as impaired mobility and a hunched physical appearance (see Supplementary Videos, V1). Kaplan-Meir survival analysis shows that whole-brain radiation provides no survival benefit (Figure 3D), as visualized by BLI (Figure 3C) and in video recordings (Supplementary Videos, V1). In contrast, mice treated with AM-101 plus radiation show a significant survival benefit and 70% of mice survived beyond radiation alone treated mice (Figure 3D). The median days to the endpoint of mice treated with AM-101 plus radiation was markedly higher than mice treated with vehicle and radiation only, 61 days versus 35 days respectively (Figure S5C). One mouse did not exhibit evidence of recurrence of an intracranial tumor by BLI (Figure 3C, at 105 days post-injection), nor at the end of the study when the brain was resected to identify macroscopic evidence of tumor. There was no significant change in body weight of AM-101 plus radiation treated mice during the entirety of the experiment, indicating no added toxicity (Figure S5D). We note, about half of all mice in each treatment group had cancerous cells in the spinal region (Figure 3C). Interestingly, mice receiving radiation localized to the brain plus AM-101 exhibited loss of tumor cells in the brain as well as spine. Further studies are required to understand the effect of AM-101 with radiation on possibly spinal or leptomeningeal metastasis. Since our heterotopic xenograft mouse experiments showed that AM-101 alone does not provide significant tumor control in immunocompromised NSG mice, we did not include an AM-101 alone treated group in the intracranial brain metastatic mouse xenograft experiments.

**Figure 3.**
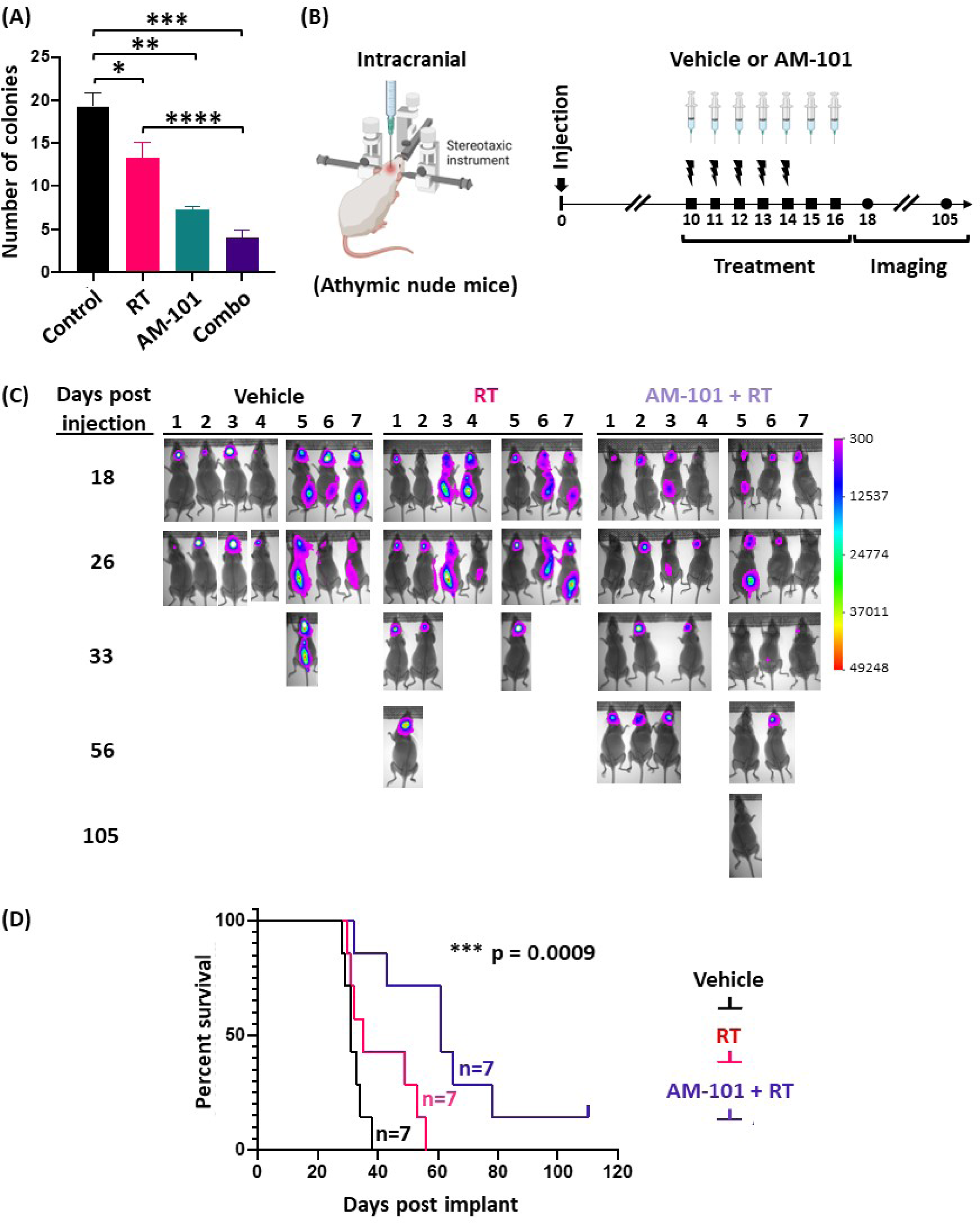
GABA(A) receptor activation increases survival of mice bearing lung brain metastatic tumors. (A) A clonogenic assay was employed to examine cytotoxicity of AM-101 combined with radiation (RT) versus AM-101 (2.5 μM) or RT (3 Gy) alone on a lung brain metastatic cell line (UW-lung-16). RT alone has a nominal effect on colony formation, while AM-101 has a two-fold greater effect. AM-101 plus RT (combo) has the most significant impact on colony formation, a four-fold greater effect than control. Control is DMSO, which is also the vehicle of AM-101. Data are represented as mean ± S.E. Student’s t test. P values between two treatment groups are: * = 0.11; ** p = 0.0004; *** p = 0.0016; **** p = 0.043. (B) Schematic of efficacy experiment in intracranial xenograft tumors generated in athymic mice. Mice received a stereotaxic intracranial injection of cells (UW-lung-16) from a brain lesion of a patient with lung cancer. Mice (n = 21) were separated into three treatment groups (n = 7 per group). Ten days post-injection, mice received: (1) vehicle, an i.p. injection of formulation; (2) RT, (2.5 Gy) to the whole mouse brain for 5 consecutive days; (3) AM-101 plus RT, i.p. injection of formulated AM-101 (5 mg/kg) for 7 consecutive days and RT (2.5 Gy dose) for 5 days to the whole mouse brain. Tumors were followed by bioluminescent imaging (BLI) over time. (C) BLI of mice with metastatic lung tumors and treated with vehicle, RT, or RT plus AM-101. Mice were imaged at indicated days post-intracranial injection. (D) Kaplan-Meier Survival Curve with p value (log-rank test) for statistical significance shown. Treatment with AM-101 plus RT shows significant survival versus mice treated with RT only.

### GABA(A) receptor activation enhances autophagic puncta and GABARAP and Nix multimerization

Having analyzed the effectiveness of AM-101 to potentiate radiation in two mouse tumor models, we examined the contribution of molecular events to the observed effect. In considering how AM-101 activation of GABA(A) receptors may mediate adenocarcinoma cell death and tumor control, several observations by us and others suggested that autophagy may be a contributing factor. First, we reported above that AM-101 activation depolarizes the mitochondrial transmembrane. It has been reported that depolarization of mitochondria serves to trigger autophagy.^38^ Second, a central protein to autophagy induction is GABARAP, which prior to elucidation of its role in autophagy, was shown to interact with GABA(A) receptors.^26^ GABARAP is also a Nix interacting factor, which has a significant role in autophagosome formation.^29^ Third, we find here that AM-101 treatment *in vitro* does not enhance cleavage of pro-caspase 3 in lung adenocarcinoma cells (Figure S4C) ever at much higher concentration that its IC_50_. Cleaved or activated pro-caspase 3 contributes to an inhibition of autophagy by catalyzing proteolysis of key autophagy-associated proteins, including Beclin-1, ATG5 and p62.^39,40^

Key proteins to the assembly of autophagy granules or puncta, a hallmark of autophagy induction, are Nix and the ATG8 sub-family associated protein LC3B and GABRAP. Importantly, Nix mediates an association between the mitochondria and GABARAP, which is associated with the GABA(A) receptor at the extracellular membrane. Subcellular distribution of autophagosomal proteins by immunofluorescence (IF) (e.g., LC3 puncta formation) is a well-established assay to monitor autophagy. It is known that lipidated LC3B (LC3B conjugated with phosphatidylethanolamine), which is present at the autophagosomal membrane, is observed as puncta within the cell cytoplasm whereas poorly lipidated LC3B shows a more diffuse staining.^40^ We therefore employed confocal IF to observe assembly of puncta in lung adenocarcinoma H1792 cells treated with AM-101 and radiation alone as well as the two combined. There is a background level of both LC3B and Nix-positive puncta in H1792 cells in the control (DMSO treated) group (Figures 4A and 4B). However, quantification of puncta reveals that AM-101 alone creates an environment for increased puncta positive for both markers (Figures 4A and 4B; see bar graphs). Similarly, radiation of cells contributes to increased puncta formation. But the most significant increase in puncta is observed when AM-101 and radiation are combined.

**Figure 4.**
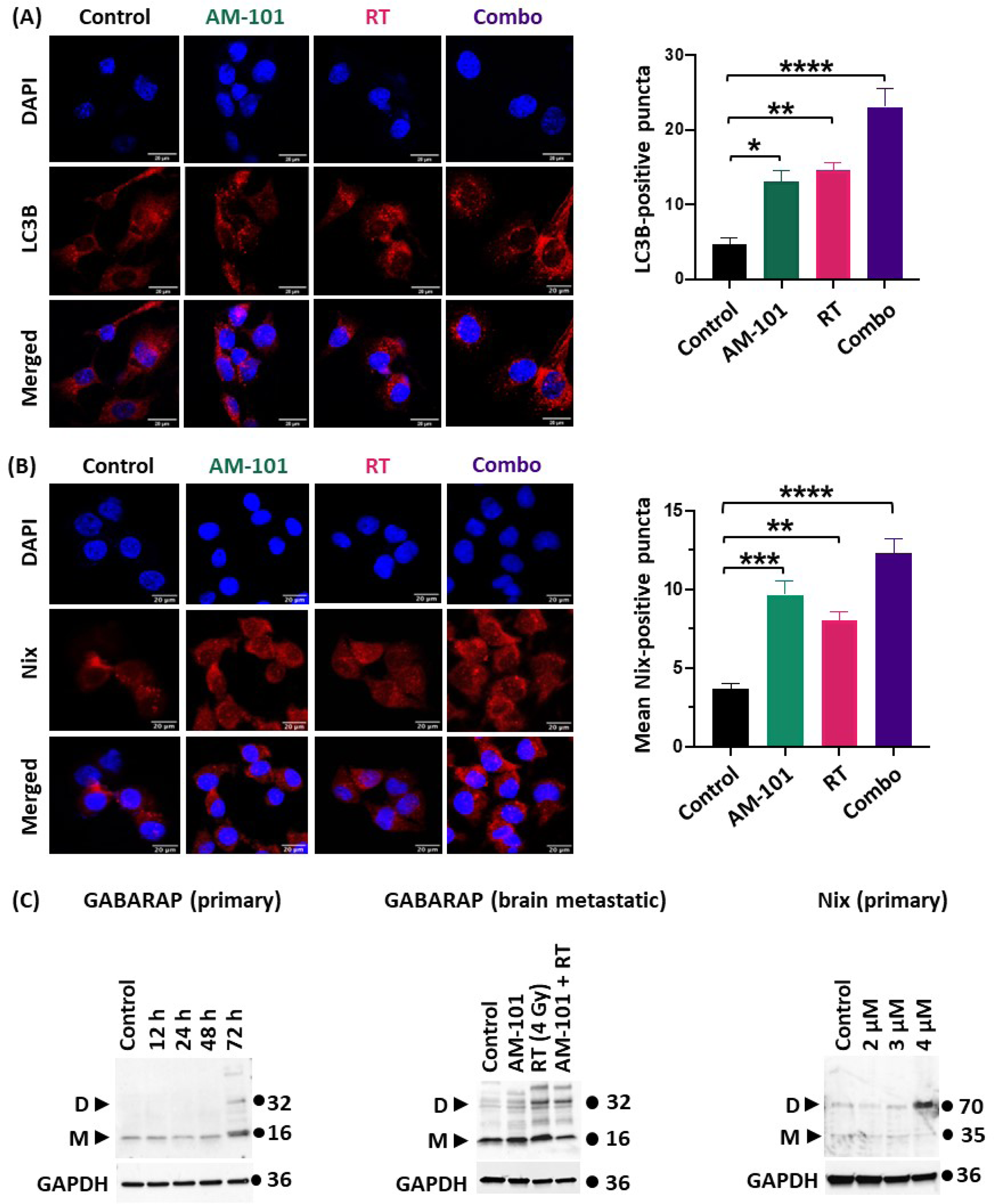
GABA(A) receptor activation enhances autophagic puncta and multimerization of GABARAP and Nix. (A) Shown are confocal immunofluorescence microscopic images of H1792 cells under various treatments: DMSO or control; AM-101; radiation (RT); and AM-101 plus RT (combo) (scale bar, 20 μm). Cells were stained for DNA with DAPI (blue fluorescent) and LC3B (left). LC3B puncta were quantified per 3 cells for each experimental group and plotted, as shown in the bar graph, where: * p = 0.0108 (Control vs AM-101); ** p = 0.0037 (Control vs RT); **** p < 0.0001 (Control vs Combo) (right), which reveals a similar effect between AM-101 versus RT but combining these two has a statistically pronounced impact on puncta formation. (B) Confocal immunofluorescence microscopic images of H1792 cells under various treatments, then stained for DAPI and Nix to identify and quantify Nix-puncta (left). Nix puncta were quantified per 3 cells for each experimental group and plotted, as shown in the bar graph, where: *** p = 0.0008 (Control vs AM-101); ** p = 0.006 (Control vs RT); **** p < 0.0001 (Control vs Combo) (right), which reveals a pronounced effect of AM-101 on puncta formation and an increase of puncta when AM-101 is combined with RT (combo treatment group). For statistical calculations one-way ANOVA was performed and followed up with Dunnett’s multiple comparisons test. (C) Immunoblot staining of SDS (4-15% gradient) gels showing the effect of AM-101 on GABARAP and Nix abundance and their oligomeric state in primary (H1792) and brain metastatic (UW-lung-16) cells. GAPDH is used as a loading control. AM-101 (3.5 μM) triggers a multimerization of GABARAP and an apparent increase in abundance at 72 hrs (left). Nix abundance is also enhanced as well as formation of dimer by AM-101 (4 μM) in a concentration-dependent manner in H1792 cells.

Having observed enhanced assembly of LC3B and Nix-positive puncta, we focused on the protein GABARAP, since Nix binds to GABARAP. In a time-course experiment, H1792 cells treated with AM-101 show a pronounced accumulation of both GABARAP and Nix at 72 hrs post-treatment. Importantly, GABARAP and Nix also dimerize at 72 hrs (Figure 4C) post treatment with AM-101. This is the time point when lung adenocarcinoma cell viability is impaired by AM-101. Furthermore, AM-101 induced Nix dimerization is dose-dependent (Figure 4C). We also observed that radiation alone as well as radiation plus AM-101 triggers GABARAP dimerization *in vitro* in brain metastatic cell line UW-lung-16, corroborating the observation in primary cells (Figure 4C). However, we did not observe an accumulation of multimers of these proteins in tumor samples. This may stem from the fact that the multimers are non-covalent complexes and most likely disrupted during tissue processing. Also, the tumors were resected at the end of the experiment when tumors have recurred.

### Enhanced levels of autophagy biomarkers in cells and tumors

Having established that GABA(A) receptor activation leads to: (1) enhanced formation of LC3B and Nix-positive puncta; and (2) GABARAP and Nix time-dependent multimerization, we turned to examining if GABA(A) receptor activation elicited an effect on the abundance and activity of other proteins key to various aspects of autophagosome assembly. Specifically, we focused on proteins ATG7, p62, and Beclin-1.

ATG7 is a Ubiquitin-activating E1-like enzyme that aids in autophagosome formation.^41–43^ AM-101 *in vitro* enhances ATG7 protein levels in both patient-derived adenocarcinoma primary (H1792) and brain metastatic (UW-lung-16) cells (Figure 5A). In the UW-lung-16 cell line this effect is time-dependent. Interestingly, of three splice variants of ATG7, the observed change in the primary cells is of a variant functionally associated with autophagy induction. Change in ATG7 levels is not only observed in H1792 cells, immunoblotting of heterotopic H1792 tumor tissue lysate shows a nominal degree of enhanced ATG7 staining in mice treated with AM-101 (Figure S6A).

**Figure 5.**
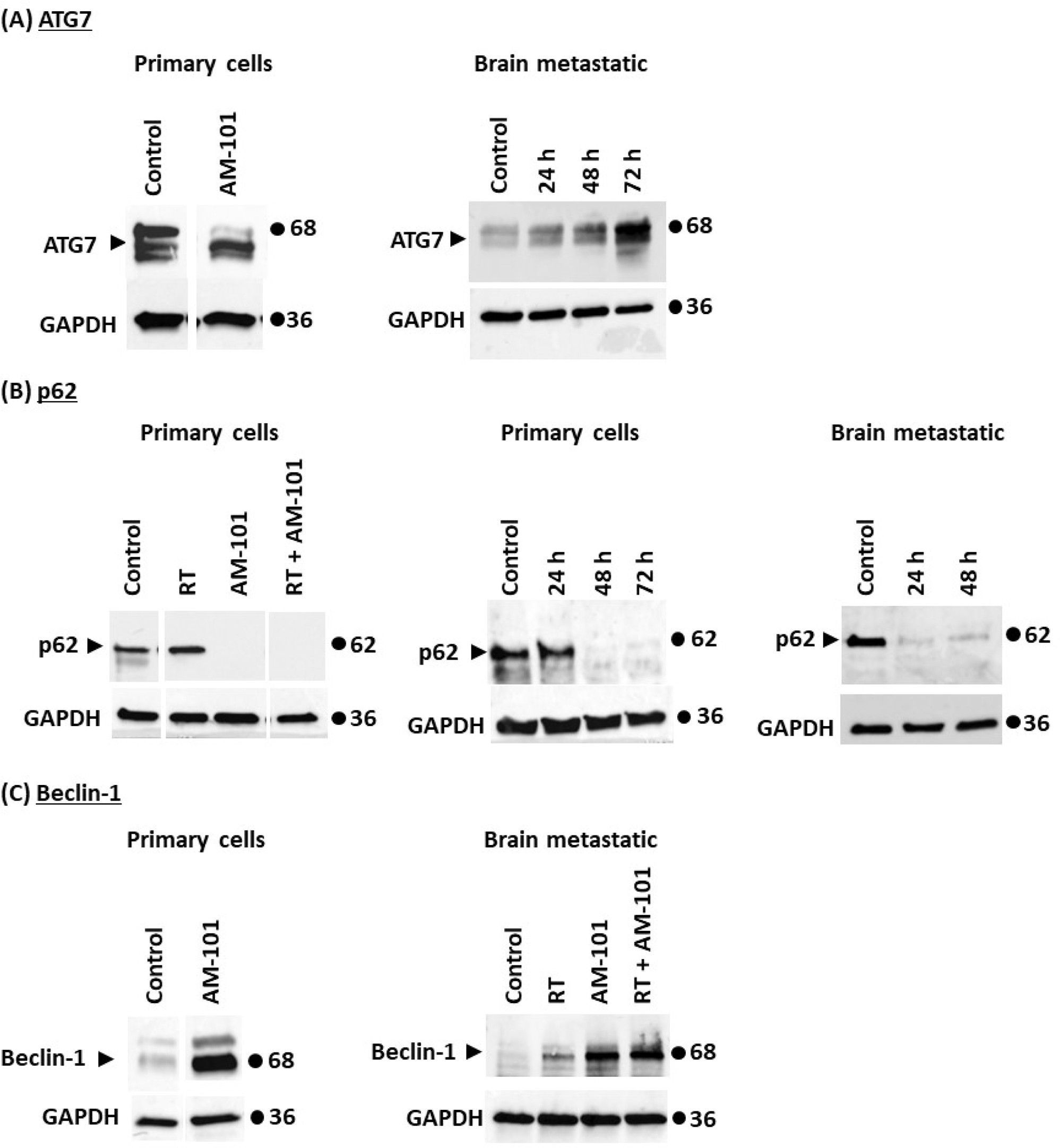
Change in abundance or utilization of autophagy biomarkers in response to GABA(A) activation. (A) ATG7 immunoblot of SDS (4-15% gradient) gels of patient-derived lung adenocarcinoma primary (H1792) and brain metastatic (UW-lung-16) cells following treatment with AM-101. Left, ATG7 abundance is enhanced of one isoform of ATG7 in primary (H1792) cells following treatment with AM-101 (3.2 mM, 48 hrs). Control: DMSO treated. Cropped gel lanes from original blots, see Figure S7B. Right, Change in abundance of ATG7 over time of brain metastatic (UW-lung-16) cells treated with AM-101 (3.2 mM) shows significant increase at 72 hrs. Control: DMSO treated. (B) p62 immunoblots of SDS (4-15% gradient) gels of lung adenocarcinoma primary (H1792) and brain metastatic (UW-lung-16) cells following treatment with AM-101. Left, Changes in abundance of p62 is assessed by immunoblotting of lysates from H1792 cells treated with AM-101, radiation (RT) (3 Gy) and a combination of RT plus AM-101. Control, DMSO. Cropped lanes from original blot, see Figure S7C. Middle, Evaluation of the time dependent utilization of p62 after AM-101 (3.2 μM) treatment of primary (H1792) cells. Right, Patient-derived brain metastatic lung adenocarcinoma cell line UW-lung-16 (right). Control, DMSO. GAPDH is used as a loading control. (C) Left, Immunoblot showing Beclin-1 protein levels in Control and AM-101 treated primary (H1792) cells (treated for 72 hours). Control, DMSO. Cropped lanes from original blot, see Figure S6D. Right, Immunoblot showing Beclin-1 levels in patient-derived lung brain metastatic UW-lung-16 cells treated with AM-101, RT) (4 Gy, or RT (4 Gy) plus AM-101.

Sequestosome-1(SQSTM1)/p62 or p62 is a ubiquitin-binding multifunctional protein which acts as a selective autophagy substrate.^44^ AM-101 alone induces a depletion of autophagy substrate protein p62, indicative of its utilization (Figure 5B). In contrast, there is no change in level of p62 in H1792 cells when exposed to radiation (3 Gy) alone. But radiation plus AM-101 depletes p62 (Figure 5B). In a time-course experiment, we found that p62 is depleted within 48 hours after AM-101 treatment of primary (H1792) and brain metastatic lung adenocarcinoma (UW-lung-16) lines (Figure 5B). In fact, the utilization of p62 appears more rapid in the metastatic cells. We also observed reduction in levels of the autophagy substrate p62 in tumor tissue lysates from the radiation only and radiation plus AM-101 treated lung brain metastatic tumors, compared to tumors from the vehicle treated group (Figure S6B).

Finally, Beclin-1 is an autophagy initiator and regulator.^45^ In culture, AM-101 enhances Beclin-1 abundance in primary (H1792) and brain metastatic (UW-lung-16) cells (Figure 5C). In contrast, relative to AM-101, we observe a marginal increase in Beclin-1 following radiation exposure (Figure 5C). Beclin-1 abundance is also enhanced in H1792 heterotopic xenograft tumors from mice treated with AM-101 or radiation alone as well as combination of the two (Figure S6C). Importantly, enhanced abundance of Beclin-1 is AM-101 dependent, as the chemically similar benzodiazepine diazepam (Valium) does not enhance Beclin-1 expression (Figure S6D). As noted above (Figures S2C and S2D), diazepam is not cytotoxic to this cancer line.

### AM-101 cytotoxicity is inhibited by abrogating the GABARAP-Nix axis

We observed that primary and brain metastatic lung cancer cells have enhanced abundance or utilization of key autophagy proteins when treated with a GABA(A) receptor activator. Our attention then turned to whether the autophagic response observed contributed to the cytotoxicity of AM-101 and the importance of GABA(A) receptor activation in triggering autophagic events. We tested autophagy inhibitor bafilomycin A1 at a low concentration (10 nm) to see if it would counteract the effect of AM-101. Bafilomycin A1 and AM-101 when combined are no more cytotoxic than each alone (Figure S7A). This suggests that they are involved in a molecular ‘tug-of-war’, opposite sides in their molecular mechanisms. We also find that bafilomycin A1 when combined with AM-101 counteracts the molecular effects reported above of AM-101, including a reduction of ATG7 abundance as well as the utilization of p62 (Figures S7B and S7C). These observations suggest that AM-101’s cytotoxic mode of action is via induction of autophagy. However, the high cytotoxic effect of bafilomycin A1 on lung adenocarcinoma cells, even at 10 nM (Figure S7A), complicates our ability to make a clear conclusion.

We therefore explored potential ways to inhibit autophagy without eliciting a cytotoxic response. We also desired an approach that had a clear target and mode of action. We employed a small peptide that had been designed to target the protein GABARAP and inhibit autophagy by blocking Nix binding.^9^ This peptide, Pen3-*ortho*, has several advantages: (1) it is highly specific for GABARAP, binding with a low nanomolar affinity; and (2) a crystal structure has been determined of Pen3-*ortho* in complex with GABARAP, thus its mode of action delineated.^9^ Mechanistically, Pen3-*ortho* binds to a site on GABARAP that is disruptive to its interaction with Nix i.e., it functions as a competitive inhibitor. We find that in contrast to bafilomycin A1, Pen3-*ortho* alone is not cytotoxic to H1792 cells or nominally at exceedingly high concentrations (Figure 6A). We then examined if Pen3-*ortho* inhibited the cytotoxic response of AM-101. Pen3-*ortho* combined with AM-101 significantly inhibits cytotoxicity of AM-101 on lung adenocarcinoma H1792 cells. This inhibitory effect is concentration dependent (Figure 6A). As expected, given its nanomolar affinity for GABARAP protein, Pen3-*ortho* does not alter GABARAP protein levels, even at concentrations well above its reported inhibitory concentration (Figure 6B). Western blotting of cells treated with Pen3-*ortho* shows significantly less Nix protein, both monomer and dimer states (Figure 6C). This supports previously published work that GABARAP interacts with and stabilizes Nix.^29^ Importantly, these experiments support GABARAP as a mediator between GABA(A) receptor function and autophagosome assembly (see Figure S7D for an illustration of this series of experiments and model based on the effect).

**Figure 6.**
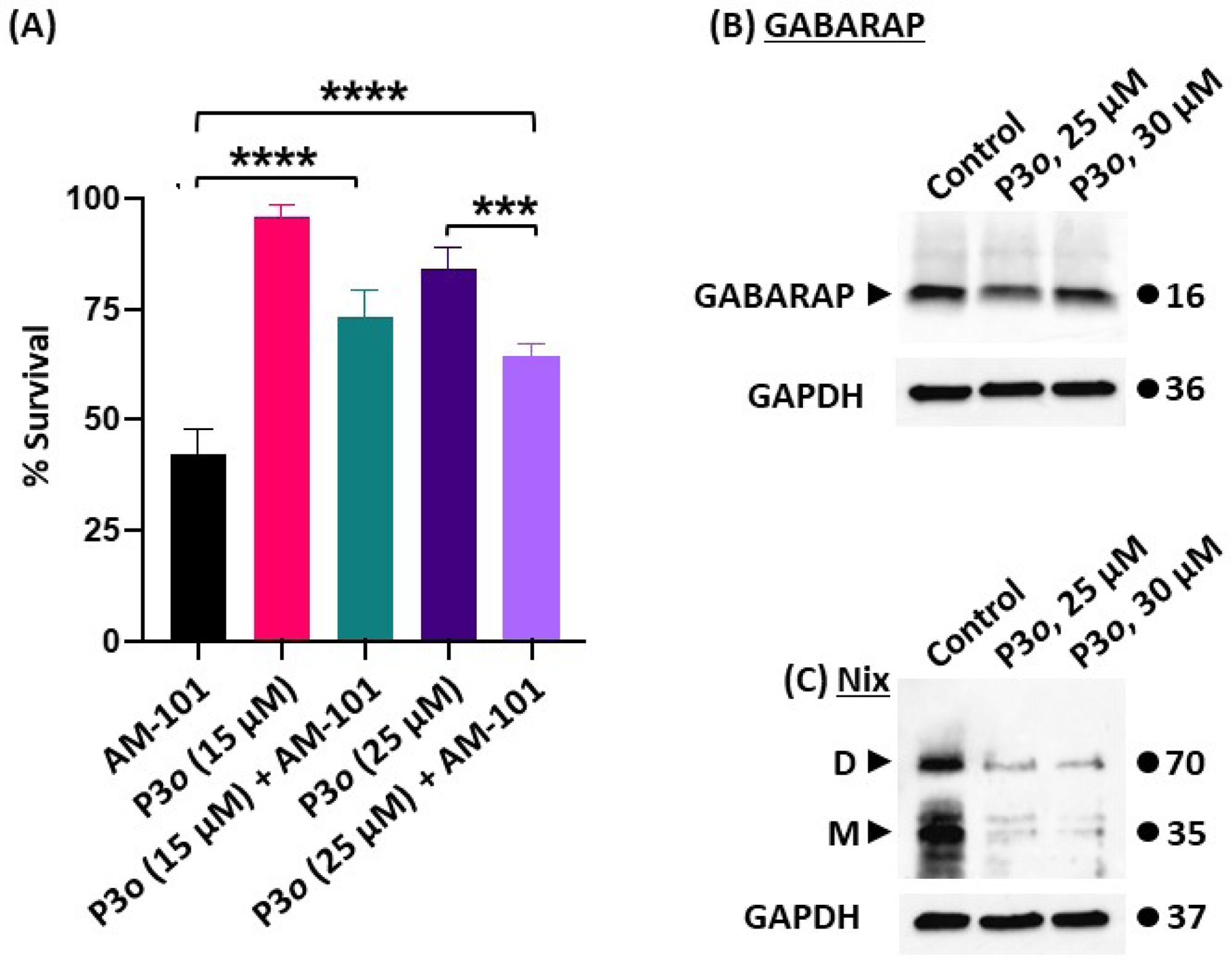
GABARAP-Nix abrogation inhibits AM-101 cytotoxicity. (A) Combined treatment of lung adenocarcinoma (H1792) cells with AM-101 plus Pen3-*ortho* (P3*o*), a stapled-peptide that binds to GABARAP and abrogates Nix binding, inhibits the cytotoxicity of AM-101. The inhibitory effect of P3*o* is enhanced with an increased concentration of the inhibitor. ****P< 0.0001 [AM-101 vs P3*o* (15 μM) + AM-101]; ****p< 0.0001 [AM-101 vs P3*o* (25 μM) + AM-101]; **** p< 0.001 [P3*o* (25 μM) vs P3*o* (25 μM) + AM-101]. One way ANOVA with multiple group comparisons were performed. (B) Treatment of H1792 cells two different concentrations of P3*o* does not impact GABARAP protein abundance, as observed by immunoblot of SDS-gel probed for GABARAP. (C) Treatment of H1792 cells with P3*o* reduces both Nix dimer and monomer protein levels, as observed by immunoblot of SDS-gel probed for Nix. D: dimer, M: monomer. GAPDH is used as a loading control.

### Contribution of γ-H2AX to radiation-sensitization

A protein which is indicative of radiation effectiveness as well as contributes to autophagy is the phosphorylated form of histone H2AX or γ-H2AX. Recently, it was reported that in embryonic stem cells, activation of GABA(A) receptors using the agonist muscimol increases γ-H2AX in response to double-strand breaks.^20,46^ Similarly, we find that AM-101 enhances γ-H2AX in lung adenocarcinoma cells (Figure S4D). Furthermore, γ-H2AX enhancement is augmented when AM-101 is combined with radiation *in vitro* (Figure S4D). AM-101 may therefore potentiate radiation via γ-H2AX. However, AM-101 enhancement of γ-H2AX is not observed in lung brain metastatic cell line UW-lung-16, although radiation alone does enhance γ-H2AX in this line (Figure S4E). Thus, while γ-H2AX may contribute to the radiation-sensitizing effect of AM-101, it appears it may not play a deciding role.

## Discussion

GABA(A) receptors are the major inhibitory neurotransmitter receptors in primates. GABA(A) receptors also have roles outside of the CNS and receptor subunits are expressed in cancer cells.^19,20^ We find in lung adenocarcinoma cells that GABA(A) receptors are functional and that their activation enhances the effect of its natural agonist, the metabolite GABA (Figure 1D). Previously, we found that activation of GABA(A) receptors in cancer cells contributes to an efflux of chloride anions.^21^ Previously and in this study, we find that GABA(A) receptor activation leads to depolarization of the mitochondrial transmembrane, consistent with a net efflux of chloride anions.^21^ Depolarization has been reported to lead to induction of autophagy.^38^ We therefore investigated changes in levels of proteins with disparate roles in autophagy, including ATG7, p62, and Beclin-1; and show changes in protein levels of these genes. We were particularly drawn to the contribution of GABARAP and Nix as proteins key to the nucleation of autophagosome assembly and ones that bridge the extracellular plasma membrane and the mitochondrial transmembrane. The GABARAP subfamily of proteins also promote autophagy by regulating the activity of kinase ULK1, whose function stabilizes autophagosome formation.^47^ In addition, phosphorylated GABARAP traffics GABA(A) receptors to the extracellular plasma membrane, binding to its γ2-subunit.^48,49^ Nix is an autophagy receptor that rests in the mitochondrial transmembrane, and complexes with GABARAP to recruit mitochondria to autophagosomes.^29^ As well as enhancing expression of GABRARAP and Nix, we find that GABA(A) receptor activation contributes to their multimerization, thus augmenting the autophagic response. We hypothesize and illustrate in our model (see Figure 7) that multimerization of GABARAP occurs commensurate with multimerization of GABA(A) receptors, and that this macromolecular assembly assumes a cytotoxic state as it drives a significant efflux of chloride anions. This overall concept is consistent with experimental analysis and theoretical modeling, which shows that multimerization of GABA(A) receptors, and in turn GABARAP, creates a strong localized distribution of charge and enhanced receptor activity.^26–28^ This would also occur and be commensurate with a stabilization and multimerization of Nix, as we observe. In this way, perturbation of ion homeostasis as sensed and regulated by GABA(A) receptors would lead to assembly of an autophagosome and potential enclosure and subsequent recycling of mitochondria.

**Figure 7.**
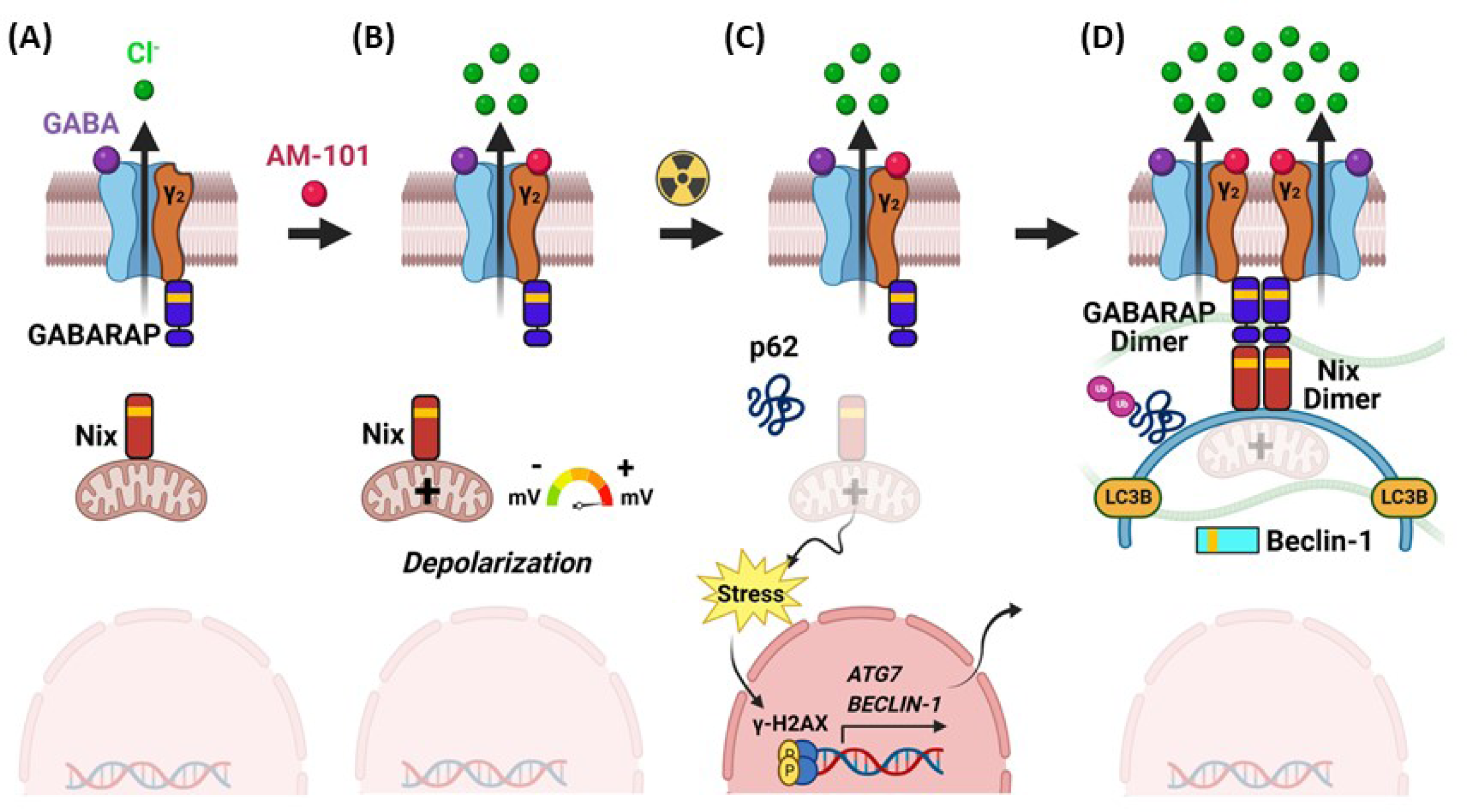
Model of GABA(A) receptor mediated autophagy. (A) NSCLC cells possess intrinsic GABA(A) receptors (chloride anion channels). (B) AM-101 activates GABA(A) receptors, which in turn depolarize the mitochondria. (C) Changes in the cancer cell by binding of AM-101 to the receptor in combination with radiation include: (*i*) enhanced expression-abundance of key genes involved in autophagy, including: *ATG7, BECLIN-1*; and (*ii*) increased phosphorylation of the histone variant H2AX to generate γ-H2AX. (D) Depolarization induces key autophagic events in synergy with radiation: (*i*) enhanced expression and dimerization of GABA(A) receptor-associated protein, GABARAP; (*ii*) stabilization and dimerization of Nix, coupling GABARAP to mitochondria; (*iii*) enhanced expression of autophagy-associated proteins Beclin-1 and ATG7; (*iv*) utilization of ubiquitin-binding protein p62. Nix dimerization increases its stability and coordinates the nucleation of autophagosome formation. In this manner, GABA(A) receptor activation induces complex multimerization activating autophagy. Over time, GABARAP multimerizes commensurate with multimerization of the GABA(A) receptor, which enhances its activity.

Interestingly, autophagy plays an important role in regulating radiation-sensitivity of disparate cancer types. In lung cancer, for example, enhanced expression of Beclin-1 overcomes radiation-resistance^50,51^ and induction of autophagy sensitizes NSCLC and glioblastoma cells to radiation.^52^ Our finding that AM-101 induces enhanced expression of Beclin-1 and reduction of the autophagy substrate p62 in NSCLC cells and tumors, supports the idea that AM-101-mediated enhancement of autophagy underpins the radiation-sensitization of tumors. Our results demonstrated that this radiation sensitization induced by AM-101 treatment translates into significant improvement in overall survival in a patient-derived xenograft mouse model of brain metastatic lung adenocarcinoma.

In conclusion, GABA(A) receptors have been a primary pharmacologic target to remediate neurological disorders for over seventy years. Here we show that pharmacologic GABA(A) receptor activation can be leveraged to treat NSCLC cancer (both in the primary and metastatic setting) in combination with radiation. AM-101 is a non-toxic, rapidly brain-penetrant^36^ radiation-sensitizer that may improve tumor control and allow for radiation dose de-intensification to reduce toxicity.

### Study limitations

We carried out an investigation to identify the effectiveness and mechanism of a GABA(A) receptor activator, AM-101, on lung adenocarcinoma brain metastasis. We employed a brain metastatic mouse model of lung adenocarcinoma which utilized intracranial implantation of human patient-derived lung adenocarcinoma brain metastatic cells and bioluminescence imaging of resulting tumor. A limitation of this model is that it may not recapitulate key cellular steps of metastasis. Specialized mouse models that employ intracardiac or intracarotid injections to generate metastatic lesions and more adaptive to magnetic resonance imaging to visualize microlesions may be more reflective. In our mechanistic study, we used a novel GABARAP binding peptide (Pen3-*ortho*) that disrupts its binding to Nix to evaluate whether AM-101 cytotoxicity is mediated through an autophagy activation mechanism. We did not confirm this result biologically by inhibition of GABARAP using a stable knockdown of GABARAP and/or Nix.

### Resource availability

#### Lead contact

Further inquiries regarding methods, reagents, or results should be directed to the lead contact, Soma Sengupta.

#### Materials availability

This study did not generate unique materials and reagents.

## Method Details

### Cell lines

H1792, H460 and H1703 were grown in RPMI (Roswell Park Memorial Institute)-1640 Medium. A459 was grown in Dulbecco’s Modified Eagle Medium (DMEM). UW-lung-2 and UW-lung-16 were grown in DMEM with glucose, L-glutamine, and sodium pyruvate.^35^ Media for all lines was supplemented with 10% (v/v) Fetal Bovine Serum (FBS) and 100 U/mL Penicillin-Streptomycin. Lines were grown at 37°C, 5% (v/v) CO_2_.

### PCR

Total RNA from confluent cells was isolated (TRIzol Reagent) and stored at −80°C. First-strand cDNA was prepared from 1 mg total RNA (ProtoScript II First Strand cDNA Synthesis Kit). RT-PCR for *GABRA5*, employed an initial denaturing step (95°C, 5 min) followed by thirty-five cycles (95°C, 30 sec; 59°C, 40 sec; 72°C, 40 sec) and a final extension (72°C, 7 min) (Platinum® PCR SuperMix High Fidelity). RT-PCR for *GABRA3* and *GAPDH* employed an initial denaturing step (95°C, 5 min) followed by thirty-two cycles (95°C, 30 sec; 57°C, 40 sec; 72°C, 40 sec) and a final extension at 72°C for 7 min.

### Immunohistochemistry

Patient lung adenocarcinoma tumor tissue (primary and matched brain metastatic) was obtained from the CLIA certified University of Cincinnati Histopathology Core Laboratory under an approved IRB. Tissue (fixed in 4% paraformaldehyde/PBS and paraffin embedded) was obtained as 5 μm sections on glass slides, which were then deparaffinized and antigen retrieved by heat induced epitope retrieval (HIER). Slides were incubated with 2.5% normal horse serum for 20 min and then with primary Ab (1:150 dilution) for 15 min at room-temperature. Polymer based HRP conjugated secondary Ab was incubated 8 min at room-temperature.

### AM-101 and Pen3-*ortho* preparation

AM-101 was synthesized as described.^53,54^ For *in vitro* and *ex vivo* studies, AM-101 was kept lyophilized at room-temperature and solubilized in dimethyl sulfoxide (DMSO, 0.125%) prior to use. For mouse studies, AM-101 was solubilized in a co-solvent formulation (propylene glycol (40%); ethanol (10%); benzyl alcohol (2%); benzoic acid (2%); sodium benzoate (2%)).^55^ AM-101 (5 mg) was dissolved in 0.1 mL of ethanol (10%) at room-temperature; then other components added with mixing, followed by 30 sec vortex mixing. Pen3-*ortho* was synthesized as described.^9^

### Electrophysiology

Functional characterization employed a Port-a-Patch single cell automated patch-clamp electrophysiology instrument (Nanion Technologies). Recording solutions (Nanion Technologies) were as follows: external solution, 140 mM NaCl; 4 mM KCl; 1 mM MgCl_2_; 2 mM CaCl_2_; 10 mM HEPES; 5 mM D-Glucose; high Ca^2+^ seal enhancer solution, 130 mM NaCl; 4 mM KCl; 1 mM MgCl_2_; 10 mM CaCl_2_; 10 mM HEPES; 5 mM D-Glucose; internal solution, 110 mM KF; 10 mM NaCl; 10 mM KCl; 10 mM EGTA; 10 mM HEPES, pH 7.2 adjusted using KOH.

To increase the current amplitude in single cell recordings, GABA and AM-101 were dissolved in a high sodium containing external solution (161 mM NaCl; 3 mM KCl; 1 mM MgCl_2_; 1.5 mM CaCl_2_; 10 mM HEPES; 6 mM D-Glucose). Whole-cell recordings were performed on cells (held at −80 mV) using a gap-free protocol under continuous perfusion of external solutions and drug application. GABA (1 μM) and AM-101 (4 μM) were applied briefly for 5 sec to record current potentiation. Data acquisition was obtained using HEKA Elektronik software. Data were low-pass filtered at 1 kHz and digitalized at 100 kHz. Data analysis was performed by computing the maximum current amplitude using Nest-O-Patch Software.

### Mitochondrial depolarization

AM-101 was diluted to 4 μM in RPMI-1640 culture media. Carbonyl cyanide 4-(trifluoromethoxy) phenylhydrazone (FCCP) was diluted to 20 μM in culture medium. Tetramethylrhodamine ethyl ester perchlorate (TMRE) was diluted to 400 nM in culture medium. H1792 cells were grown in culture to 75 – 90% confluency. Cells (5×10^5^ cells/mL) were harvested and resuspended in media. Cell suspension (200 μL) was dispensed and AM-101 or FCCP (200 μL) added to final concentrations of 2 μM and 10 μM, respectively. Solution was briefly vortexed and incubated for 10 minutes. TMRE (40 μL of 400 nM stock) was added (final concentration 10 nM), vortexed, incubated for 2 minutes and sample reading was acquired using a BD LSR Fortessa. Data was analyzed using Flowjo v10 software.

### Lung cancer-chips

Human Alveolar Epithelial Cells (HPAECs) were cultured on a Matrigel/collagen coated T25 flask for 4-7 days in serum-free Small Airway Epithelial Cell Growth Medium (SAGM) and used without further passaging. Human Lung-Microvascular Endothelial Cells (HMVEC-L) were grown on a collagen I coated flask (T75) in microvascular cell culture medium and used between passage 4-5.

Lung-chips are composed of transparent material made of a tetrafluoroethylene-propylene (FEPM) elastomer containing two parallel microchannels separated by a thin collagen vitrigel membrane, previously described, and characterized.^56^ In contrast to previous organ-on-Chip models of the lung,^57–59^ we used a Polydimethylsiloxane (PDMS)-free design. The chips were a kind gift of Naoki Matsuoka. Before experiment, both microchannels were chemically functionalized using 3-aminopropyl-trimethoxysilane (APTMES) to covalently bind extracellular matrix (ECM) proteins before seeding human cells, as previously described.^60^ The apical or epithelial channel (1 × 1 mm) was coated with a mix of human placenta collagen IV (200 μg/mL), human placenta laminin (10 μg/mL), and human plasma fibronectin (30 μg/mL), all resuspended in Hanks’ Balanced Salt Solution (HBSS). The bottom or vascular channel (200 μm × 1 mm) was coated with human placenta collagen type IV (200 μg/mL) and human plasma fibronectin (30 μg/mL). Chips were incubated overnight at 37°C to complete the surface coating. The next day, HPAECs were seeded in the epithelial channel at a density of ∼ 0.8 × 10^6^ cells/mL in SAGM and chips were incubated at 37°C. Three hrs after cell seeding, both epithelial and vascular channels were rinsed 2X with 100 μL SAGM to remove unattached cells. Medium was then replaced with SAGM supplemented with keratinocyte growth factor (10 ng/mL), isobutylmethylxanthine (100 μM), 8-bromo-cyclicAMP (100 μM), and dexamethasone (200 μM). The medium in the epithelial channel (SAGM + KIAD) was replaced once a day for 6 days. Three days after epithelial cell seeding, the vascular compartment was seeded with HMVEC-L at a density of ∼ 8×10^6^ cells/mL in microvascular cell culture medium (EGM-2 MV), chips were then incubated at 37°C. One hour after endothelial cell seeding, the vascular channel was rinsed 2X with cell culture medium to remove free cells. The vascular medium was then refreshed once a day with EGM-2 MV for 3 days. One day after endothelial cell seeding, H1792-GFP cancer cells were seeded on the epithelial compartment at a density of ∼ 3.5×10^5^ cells/mL, as previously described.^58^ The epithelial channel was rinsed with fresh cell culture medium 3-5 hrs after seeding of cancer cells and air-liquid interface (ALI) was established the next day via pipetting 50 μL of air in the epithelial channel. Chips were then connected to flow using a syringe pump and perfused with a modified endothelial cell culture medium (ALI medium) made with Medium 199 supplemented with 2% HyClone FetalClone II Serum; 10 ng/mL Epidermal Growth Factor, 10 ng/mL Keratinocyte Growth Factor, 0.25 ng/mL Vascular Endothelial Growth, 1 μg/mL Hydrocortisone, 1 U/mL Heparin, 50 μM 8-Bromoadenosine 3′,5′-cyclic monophosphate, GlutaMAX, 20 nM Dexamethasone, and Penicillin-Streptomycin.

### Immunoblotting

Tumor tissue (heterotopic xenograft or brain) was harvested, minced on ice, and lysed in ice cold RIPA buffer supplemented with protease and phosphatase inhibitor cocktail. DNA was sheared by sonication. Lysates were kept on ice for 30 min, centrifuged 10 min (13,500 × *g*, 4°C), and protein concentration of supernatant determined by a Bradford assay. Lysates were mixed 1:1 with 2X Laemmli sample buffer containing β-mercaptoethanol and heated 5 min, 95°C. Protein (∼ 30 μg) was resolved by SDS-PAGE using 4-20% gradient polyacrylamide gels, then transferred to nitrocellulose membranes for 2 hr at 100 V in tris-glycine transfer buffer containing 20% methanol. Membranes were blocked at room-temperature in 5% 1x TBST blocking buffer (TBS with 0.1% Tween-20 and 5% non-fat dry milk) for 1 hr with gentle agitation, followed by overnight incubation with primary antibody and gentle shaking at 4°C. The primary antibodies were diluted as follows: GABRA5 (1:1000); β-actin (1:1000); GAPDH (1:1000); GABARAP (1:1000; ATG7 (1:1000); NIX (1:1000; p62/SQSTM1 (1:1000 in 5% milk in TBST). Immunodetection was performed with anti-rabbit horseradish-peroxidase-conjugated secondary antibody (1:10000). Post-primary Ab incubation, membranes were washed (3X for 10 min in TBST at room-temperature), probed with rabbit HRP tagged secondary antibody (1:3000), and processed for chemiluminescence detection using ECL kit. Chemiluminescence images were acquired using ChemiDoc Touch Imaging System.

### Microscopy

Cells were seeded on sterile glass cover slips and grown overnight at 37°C. Cells were rinsed with cold PBS and fixed in 4% paraformaldehyde (30 min, room-temperature). Fixed cells were rinsed in PBS (3X 5 min at room-temperature), and permeabilized (15 min in PBS containing 0.1% Triton X-100). Cells were washed in PBS (3X 5 min at room-temperature) and incubated with blocking buffer (PBS containing 0.3% Triton X-100 and 3% BSA) for 1 hr at room-temperature with gentle shaking. Blocking solution was aspirated, cells washed with ice-cold PBS, and incubated overnight with primary Ab (LC3B or Nix depending on the experiment) diluted 1:200 in sterile 0.5% BSA in PBS at 4°C on the cover slip placed on a glass slide kept inside a humidified 10 cm dish.^61^ Cells were washed (3X for 5 min in PBS) and incubated with fluorophore conjugated secondary Ab goat anti-rabbit Alexa Fluor 594 in 2% normal donkey serum at room-temperature for 1 hr in dark. Cells were washed (3X for 5 min in PBS) under low light and mounted on a glass slide in Vectashield Antifade Mounting Medium with DAPI. The following Ab’s were used (1:200 dilution): rabbit anti-human LC3B and rabbit Nix. Cells were also prepared as negative control where the primary antibody was omitted. Slides were imaged on a Zeiss LSM 710 laser scanning confocal microscope and analyzed using NIH Fiji ImageJ2.^62^

### Viability assays

For the clonogenic assay, a fixed number of cells were seeded (120 cells per well, H1792; 150 cells per well, UW-lung-16), grown overnight in humidified conditions at 37°C, and treated with AM-101 or DSMO for 1 hr before irradiation. Each treatment group had four replicates. Once colonies averaged 40 or more cells (14-18 days in control wells), media was removed, and cells washed with sterile PBS. Cells were fixed for 20 min with methanol-acetone (3:1 by volume) and stained with 1% (w/v) crystal violet in methanol for 20 minutes. Stained colonies were gently washed with distilled water, dried, and imaged. Colonies of 40 or more cells counted. Cells in radiation groups were irradiated one day after seeding with a single fraction X-ray source (Xstrahl Ltd.) at room-temperature. XenX irradiator has a collimator that delivers a uniform dose and a PA beam (posterior-anterior) focused on top of wells. In the Control group, media was replaced by fresh media after 72 hr. Colonies were allowed to grow in each of the experimental groups for 14 days (H1792 cells) and 18 days (UW-lung-16 cells) following irradiation. Media was refreshed after every 4 days and AM-101 treatment was done for 72 hours. Media was removed and colonies washed with PBS and fixed staining. The clonogenic survival curve for each condition was fitted to a linear quadratic model according to least squares fit, weighted to minimize the relative distances squared, and was compared using the extra sum-of-squares F test. Each point represents the mean surviving fraction calculated from four replicates and error bars represent the standard error (SE). Data is represented as mean ± SE of colonies using bar graphs and was analyzed using GraphPad Prism 8. p-values were calculated using one-way ANOVA.

Cell proliferation viability studies employed the Cell Titer 96^®^ Aqueous One Solution Assay.^32^ IC_50_ values were determined using the ‘[Inhibitor] versus normalized response’ nonlinear regression function and the log of inhibitor concentration in GraphPad Prism 8. For cell survival assays with Pen3-*ortho* stapled-peptide and AM-101, H1792 cells were plated with phenol-red free RPMI-1640 media (2500 cells per well in 100 μL media) and grown overnight. Next day media was removed, fresh 100 μL phenol-red free RPMI media added, and the following treatment groups setup: (1) Pen3-*ortho* group (15 μM); (2) Pen3-*ortho* (25 μM); (3) AM-101 group (3 μM); (4) combination group at AM-101 (3 μM) and Pen3-*ortho* (15 μM); (5) combination group at AM-101 (3 μM) and Pen3-*ortho* (25 μM); (6) DMSO control with no drug added; (7) control with no cells added, i.e. media alone. Cells were then incubated at 37°C, 5% CO_2_ for 48 hrs. Following, 20 μL of diluted MTS reagent was added, cells incubated at 37°C for 1 hr, and absorbance (490 nm) measured using a microplate reader. The mean of the absorbance of media only samples was subtracted from control and test group samples. The percentage of inhibition in each group was calculated by the formula (C-T)/C x 100%; where: ‘C’, is the mean absorbance reading for Control group; ‘T, is the mean absorbance reading for each treated group. The percentage of survival was calculated (GraphPad Prism 8.0.1) by subtracting percentage of inhibition of each group from 100% and expressed as mean ± SEM. Student’s t test (paired) for two groups were used for statistical comparison. A p < 0.05 is considered significant.

### Mouse experiments

Mice were housed in pathogen-free rooms and clinical health evaluated weekly by veterinary staff (University of Cincinnati LAMS). All animal studies were conducted in accordance with IACUC approval (University of Cincinnati). For subcutaneous xenograft tumor growth delay experiments, NOD-SCID gamma (NSG) mice were used. For intracranial xenograft tumor experiments, athymic nude mice were used. For mice receiving radiation, tumors were irradiated using an XenX small animal irradiator.

For heterotopic xenograft experiments, H1792 cells (0.5 million) grown in RPMI medium were washed in cold PBS, mixed with Matrigel (25%), and injected subcutaneously into left and right flanks above the hind limbs of 6- to 8-week old female mice. When subcutaneous tumors were palpable (35 days; ∼ 100 mm^3^) the following treatments groups were initiated: (1) vehicle; (2) radiation; (3) AM-101 ± radiation; and (4) docetaxel ± radiation. In vehicle mice, vehicle was injected i.p for 7 days. In radiation mice, left-side flank tumors were irradiated with a single 5 Gy dose. In radiation plus drug mice, AM-101 (2.5 mg/kg body weight) or docetaxel (8 mg/kg body weight) was injected i.p. 1 hr prior to irradiation with a single 5 Gy dose. AM-101 or docetaxel was subsequently injected i.p. for 7 days. Following treatment, tumor volume and body measurements were taken for growth delay studies. Mice were euthanized using CO_2_ euthanasia when the experimental endpoints according to the animal care guidelines were reached, subcutaneous tumors were excised and weighed and preserved at appropriate conditions for protein and IHC studies. The day of tumor cell implantation in mice was assigned as day zero in the timeline of the experiment. Mouse tumors were measured by Vernier calipers. Tumor volume was calculated using the formula: (π/6) x (*l* x *h*^2^), where: *l* and *h* are large diameter and small diameter of the tumor taken perpendicular to each other.

For intracranial experiments, UW-lung-16 cells stably transduced with luciferase were harvested from culture and washed in sterile PBS and suspended in cold sterile PBS. Viable cell numbers were calculated using Trypan blue staining and adjusted for intracranial implantation. Mice eyes were protected using an Artificial Tears solution. Mice were put under isoflurane anesthesia using an inhalation anesthesia machine (VetEquip). Analgesic buprenorphine was administered subcutaneously to each mouse before starting the procedure. The skin on the skull of mouse to be injected was disinfected using isopropanol and chlorohexidine gluconate and a small incision was made on the skin to expose the skull. From the bregma, the syringe was moved to the appropriate AP and lateral coordinates, and a small hole was drilled using a 26-gauze needle. Using a Hamilton syringe, 2×10^5^ cells suspended in 3 mL sterile PBS were implanted by stereotactic injection into the right striatum of each 6-week old athymic nude mouse with the following stereotactic landmarks: 2 mm right lateral and 0.5 mm frontal to the bregma at 3 mm depth. The stereotactic injection occurred over 5-6 min. Following stereotactic injection, bone wax was applied at the hole in the skull and scalp closed with sterile staples. Mice were kept on a heating pad during the procedure and to recover from anesthesia. The calculated median survival of mice with implanted brain metastatic tumors and receiving no treatment was 31 days after implantation. Successful tumor engraftment was ultimately confirmed by bioluminescence imaging (Druker) 8 days after implantation. Eight days after implantation, animals were randomly assigned to three experimental groups: (1) Vehicle only; (2) radiation plus vehicle; (3) AM-101 plus radiation (n = 7 mice per group). AM-101 was administered at a dose of 5 mg/kg per mouse by i.p. injection daily for 7 days. Whole brain radiation was administered 2.5 Gy daily fraction for 5 days using XenX small animal irradiator and a supplied mouse gantry. AM-101 was given 1 hr before radiation treatment. The day of tumor implantation was assigned as day zero. Mouse brain tumor growth and spinal lesion(s) (if any) were evaluated by luciferase based BLI on days 8, 18, 26, 33, and 56 and 105. For luciferase imaging each mice received intraperitoneal injections of freshly prepared sterile D-Luciferin, potassium salt solution (15 mg/mL) in 0.9% saline solution. Each mice received D-luciferin injection based on their body weight (10 µl of stock solution per 10 gram of body weight). Mice were imaged 10 minutes after D-luciferin injections. The body weight of mice was also measured over time and mice were regularly monitored for symptoms. Mice were euthanized when they reached experimental endpoint. Brain tumor tissue from cerebral cortex was dissected and processed for analysis.

### Data analysis and statistics

Data for cell survival experiments is expressed as mean ± SEM. Data analysis employed GraphPad Prism 8.0.1 software. Statistical differences were determined by two-tailed unpaired Student’s t-test for two groups. To adjust for the multiplicity, while comparing two groups at a time, Bonferroni correction was included, and the p-value threshold adjusted accordingly. One-way ANOVA with Turkey’s post-tests for multiple group comparisons was performed with the assumption of Gaussian distribution of residuals. A p < 0.05 is considered significant. All figures were assembled with Microsoft Office Professional Plus 2019.

## Data and code availability

- Original data are available from the lead contact upon request.
- This paper does not report original code.
- Any additional information required to reanalyze the data reported in this paper is available from the lead contact upon request.

## Acknowledgments

We thank Dr. Subrahmanya Vallabhapurapu and Professor Xiaoyang Qi of the Tumor Modeling Program of the University of Cincinnati Brain Tumor Center for help with intracranial injections. We acknowledge the University of Cincinnati Histopathology Core Laboratory for providing IHC services and the University of Cincinnati Biorepository for providing human lung adenocarcinoma and patient-matched lung adenocarcinoma brain metastasis tumor samples. We also thank Sharon Wang and Lisa Lemen from the University of Cincinnati preclinical imaging core for their help with bioluminescence image acquisition and mice heterotopic and whole-brain irradiation. We thank the following sources of financial support: R01 AA029023/AA/NIAAA NIH HHS, R01 DA043204/DA/NIDA NIH HHS, R01 DA054177/DA/NIDA NIH HHS, and CHE-1625735/NSF Division of Chemistry to J.M.C.; GM148407 from the National Institute of Health (NIH) to J.A.K.; R01CA273586 from the NIH to C.W.; Faculty Scholars Research Award at the University of Cincinnati to R.B.; Thomas E. & Pamela M. Mischell Family Foundation, the Harold C. Schott Foundation funding of the Harold C. Schott Endowed Chair, University of Cincinnati College of Medicine, and the American Brain Tumor Association Discovery Grant Fully supported by an Anonymous Family Foundation to S.S. We also acknowledge the University of Wisconsin-Milwaukee Shimadzu Laboratory for Advanced and Applied Analytical Chemistry; the Milwaukee Institute of Drug Discovery and the University of Wisconsin-Milwaukee Research Foundation; and Naoki Matsuoka at AGC Inc., Tokyo Japan, for providing non absorptive microfluidic Chips (DOI: 10.1021/acsomega.1c03719).

## Author contributions

D.B., D.A.P.K., S.S. were involved in conceptualization and designing the experiments. D.B. and D.A.P.K. were responsible for the original manuscript draft writing. S.S. and D.A.P.K. conducted the overall supervision and funding acquisition. D.B., R.B., D.K.T., V.S.G., D.I. performed experiments and data analysis. A.K. and P.B.D. were responsible for developing an AM-101 formulation. H.B. and J.A.K. developed the stapled-peptide. T.A., S.R., and J.M.C. developed and provided AM-101. N.H. supervised overall statistical analysis. L.K., K.W., and C.W. were involved in reviewing, and editing. K.W. and D.I. provided guidance and commentary on radiation experiments. A.M.B. provided patient-derived brain metastatic cell lines, as well as review and editing. M.M. conducted gene expression analysis using TCGA data. All authors approved the submission.

