## Supplemental Figures for "GABA(A) receptor activation drives GABARAP-Nix mediated autophagy to radiation-sensitize primary and brain-metastatic lung adenocarcinoma tumors"

### Supplementary Figures

#### Figure S1 GABA(A) receptor subunit gene expression in lung cancers

Heatmaps of GABA(A) receptor mRNA expression analysis from RNA-seq datasets of patient samples from pre-processed TCGA lung squamous cell carcinoma (A) and adenocarcinoma (B) primary tumors. The pre-processed TCGA lung squamous cell carcinoma (SC)[1] and lung adenocarcinoma (AC)[2] RNA-seq datasets were downloaded from the integrative LINCS (iLINCS) portal[3] and filtered to remove metastatic and recurring tumor samples. The expression profiles of GABA(A) genes were clustered and heatmaps were created using Morpheus (https://software.broadinstitute.org/morpheus/). (C) GABRA5 and (D) GABRA3 mRNA levels were assessed by RT-PCR in a panel of human NSCLC cell lines. GAPDH mRNA was amplified and used as an endogenous control. (E) IHC of GABRA3 stained sections of human primary lung adenocarcinoma tissue section (left) and patient matched tissue section of lung brain metastatic adenocarcinoma (right).

#### Figure S2 Effect of AM-101 and diazepam on survival of non-small cell lung cancer cells

Log transformed curves of AM-101 incubated with (A) lung adenocarcinoma (H1792) and (B) lung brain metastatic (UW-lung-16-GFP-Luc) cells. (C) Chemical structures of diazepam (left) and AM-101 (middle) showing the ethinyl bond in place of a chloride in diazepam (Valium) at position 7 in the 1,4-benzodiazepine ring system. Log transformed curves from *in vitro* MTS assay with diazepam on H1792 cells (right). Due to lack of a cytotoxic effect, an $IC_{50}$ value is not reported. (D) Effect of treatment of AM-101 (3.0 μM), diazepam (20 μM), or Control, DMSO, on H1792 cell viability, as analyzed by bright field microscopy.

**Figure S3 Human-relevant *ex vivo* 'lung-on-chip'**

(A) Immunofluorescent staining of the Lung-on-Chip's epithelial compartment co-cultured with H1792-GFP cells (green) reveals actin filaments (red, phalloidin) and nuclei (blue, HOECHST) localization. (B) The vascular compartment exhibits staining for PECAM1 (magenta, endothelial cell marker), actin filaments (cyan, phalloidin), and nuclei (grey, HOECHST). (C) H1792-GFP cells grow and spread through the epithelial compartment of the Lung-on-Chip over time. (D) Results of image analysis and quantitation of the GFP signal spread over time expressed as percentage of area covered by the GFP signal. (E) H1792-GFP cells undergo cell death in response to docetaxel, leading to (F) a reduction in the number of GFP-positive cells.

**Figure S4 Effect of AM-101 and radiation on clonogenicity of lung adenocarcinoma cells, Caspase 3 cleavage, and γ-H2AX induction**

(A) Effect of AM-101 and radiation (RT) on clonogenicity of lung adenocarcinoma (H1792) cells. RT dose, 3 Gy; AM-101, 2.5 μM. (B) Effect of AM-101 and radiation on clonogenicity of metastatic (UW-lung-16) cells. RT dose, 3 Gy; AM-101, 2.5 μM. (C) Effect of AM-101 treatment on caspase-3 cleavage in lung adenocarcinoma (H1792) cells analyzed by immunoblot of SDS (4-15% gradient) gel. Various concentrations of AM-101 were incubated with H1792 cells for 48 hrs. Lysate was then probed with Caspase-3 antibody (CST). Control in this experiment was DMSO. GAPDH is used as a loading control. (D) Immunoblot of DNA damage marker γ-H2AX in lysates of primary (H1792) cells treated with either RT (4 Gy), AM-101 (3 μM), or RT plus AM-101. Lysates were collected 20 hrs after treatment. GAPDH is used as a loading control. (E) Immunoblot of γ-H2AX from lysates from metastatic (UW-lung-16) cells treated with either RT (4 Gy), AM-101 (3 μM), or RT plus AM-101. Lysates were collected 20 hours after treatment. GAPDH is used as a loading control.

**Figure S5 Mouse xenograft experiments**

(A) Weights of excised subcutaneous H1792 xenograft tumors in mice from different treatment groups at their experimental endpoints are represented in a scatterplot graph. Data are represented as mean ± S.E. Student's t-test was performed to compare the means of two respective groups and to adjust for the multiplicity Bonferroni's correction. The p-values between each pair of groups compared are indicated. Due to the Bonferroni correction and as we did four different comparisons, a value of $p < 0.0125$ is considered significant. (B) Bar graphs showing median days to the endpoint in different treatment groups of mice bearing UW-lung-16 intracranial orthotopic xenograft tumors. (C) Flux intensities of UW-lung-16 implanted brain metastatic tumors at different time points of mice treated with radiation (RT) or RT plus AM-101. (D) Mean body weights of UW-lung-16 intracranial brain metastatic tumor bearing mice from three treatment groups: vehicle or control; RT; or RT plus AM-101. Body weight of mice was recorded over time.

**Figure S6 Change in abundance of autophagy biomarkers in tumors in response to GABA(A) activation**

(A) Immunoblot of ATG7 in lysates from H1792 tumors from different treatment groups. Radiation (RT), 5 Gy; L, left flank tumor; R, Right flank tumor. (B) Immunoblot of intracranial tumor tissue lysate. RT shows reduced level of p62, indicative of its utilization. Combined treatment of RT plus AM-101 shows a greater utilization (based on band intensity). (C)

Immunoblots of primary (H1792) and metastatic (UW-lung-16) tumors following treatment with AM-101 plus RT. Left, Beclin-1 abundance is enhanced in H1792 tumors by RT and AM-101 alone. Combined treatment does not appear to enhance Beclin-1 in tumors relative to AM-101 alone. RT dose: 5 Gy, L: left flank tumor; R: Right flank tumor. Control, vehicle treated. Right, Immunoblot to detect Beclin-1 abundance in intracranial (UW-lung-16) tumor tissue harvested from treatment groups: vehicle; RT; RT plus AM-101. (D) Immunoblot of Beclin-1 showing a time-dependent appearance following treatment with AM-101 or diazepam (Valium) of primary (H1792) cells. GAPDH is used as a loading control.

**Figure S7 Inhibition of AM-101 autophagy by bafilomycin A1 and mechanism of action of GABARP binding stapled peptide Pen3-*ortho***

Bar graphs showing results of an MTS assay (performed 48 hrs post-treatment) demonstrating the effect of combining autophagy inhibitor bafilomycin A1 (10 nM for 5 hrs incubation) with AM-101 on human H1792 NSCLC cell survival. (B) Change in expression of ATG7 in human H1792 NSCLC cells treated *in vitro* either with AM-101 and/or bafilomycin A1. (C) Change in expression of p62 in H1792 cells treated with AM-101, bafilomycin A1 (50 nM), radiation (RT, 3 Gy), and combination of bafilomycin A1, AM-101 and/or RT. Control, DMSO. GAPDH is used as a loading control. (D) Model of how Pen3-*ortho* may inhibit the cytotoxic effect of AM-101. (1) Pen3-*ortho* binds GABARAP in the same pocket where Nix associates with GABARAP, thus exhibiting a competitive inhibition mechanism. (2) AM-101 treatment promotes GABARAP and Nix interaction which triggers autophagosome membrane formation leading to autophagy. (3) Pen3-*ortho* treatment with AM-101 competitively inhibits binding of

GABARAP with Nix and reduces Nix monomer and dimer, thereby attenuating the pro-autophagy effect of AM-101.

**Supplementary Video V1 Videography of mouse intracranial experiments**

Videos of mice that received: vehicle; vehicle plus radiation (RT); or AM-101 plus RT. Videos highlight health of mice and symptoms over time post-treatment. Vehicle only mice videos: Vehicle Day 26 Mice 1,2,3,4; Vehicle plus RT mice videos: Vehicle + RT Day 30 Mice 1,3; Vehicle + RT Day 32 Mice 5,6,7; AM-101 + RT mice videos: AM-101 + RT Day 38 Mice 1,2,3; AM-101 + RT Day 38 Mice 4,6,7; AM-101 + RT Day Mice 53 2,3; AM-101 + RT Day 53 Mice 4,6,7; AM-101 + RT Day 78 Mice 4,6; AM-101 + RT Day 85 Mice 4. Numbers for mice correspond to the numbers for mice shown in Figure 3B.

Figure S1

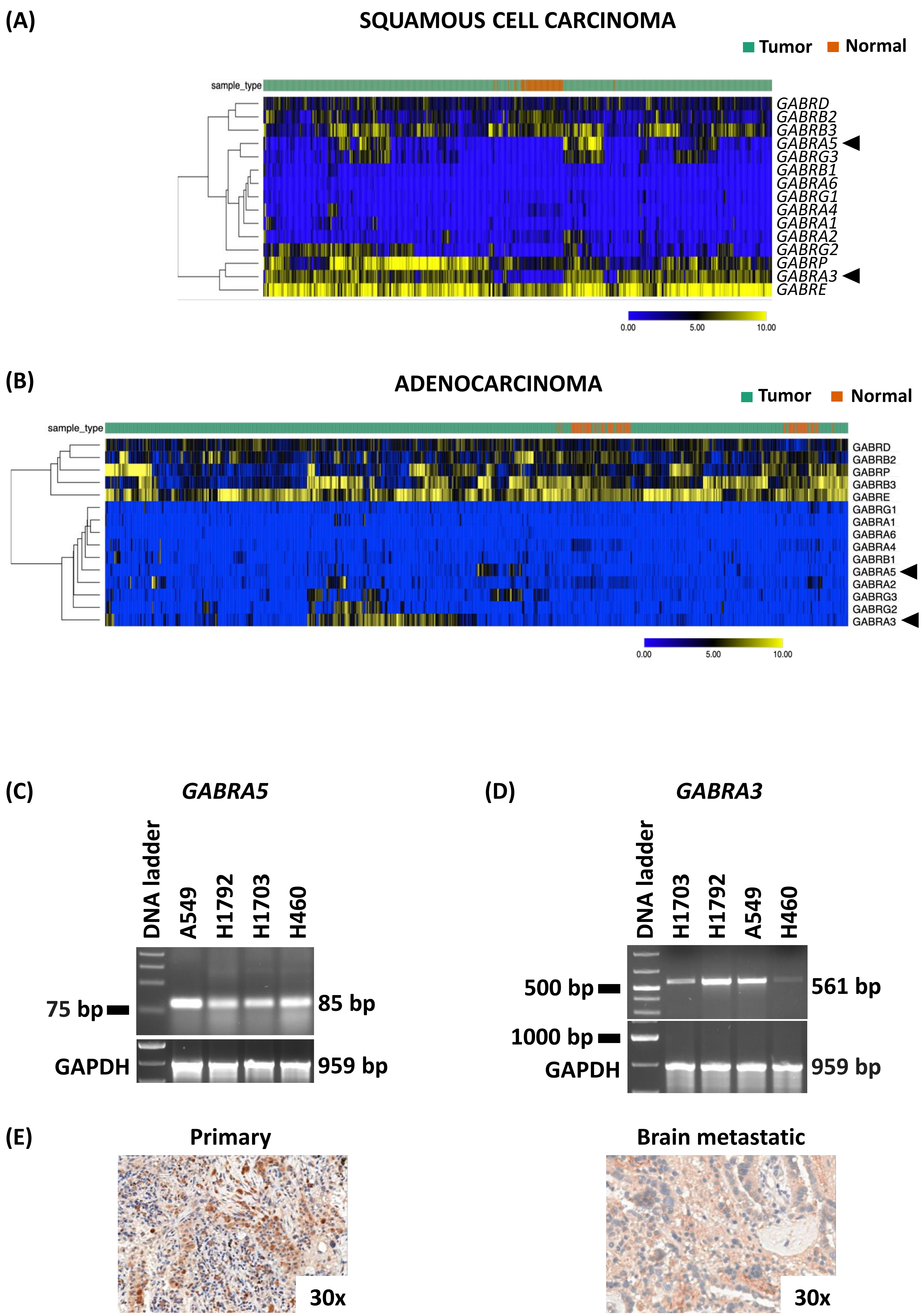

(A)
SQUAMOUS CELL CARCINOMA
Tumor
Normal
sample_type
GABRD
GABRB2
GABRB3
GABRA5
GABRG3
GABRB1
GABRA6
GABRG1
GABRA4
GABRA1
GABRA2
GABRG2
GABRP
GABRA3
GABRE
0.00
5.00
10.00
(B)
ADENOCARCINOMA
Tumor
Normal
sample_type
GABRD
GABRB2
GABRP
GABRB3
GABRE
GABRG1
GABRA1
GABRA6
GABRA4
GABRB1
GABRA5
GABRA2
GABRG3
GABRG2
GABRA3
0.00
5.00
10.00
(C)
GABRA5
DNA ladder
A549
H1792
H1703
H460
75 bp
85 bp
GAPDH
959 bp
(D)
GABRA3
DNA ladder
H1703
H1792
A549
H460
500 bp
561 bp
1000 bp
GAPDH
959 bp
(E)
Primary
30x
Brain metastatic
30x

### Figure S2

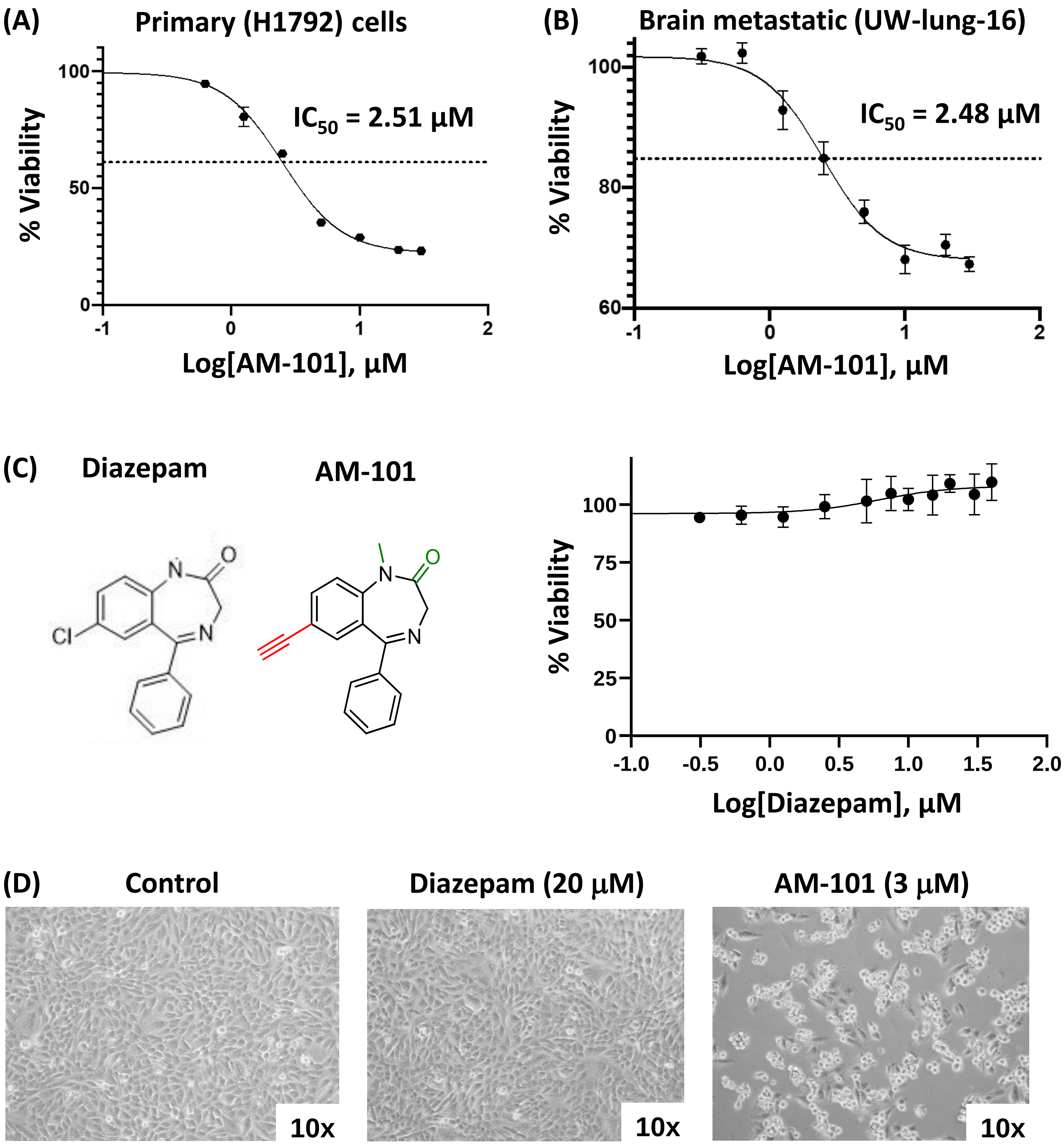

(A)
Primary (H1792) cells
IC50 = 2.51 µM
% Viability
Log[AM-101], µM
(B)
Brain metastatic (UW-lung-16)
IC50 = 2.48 µM
% Viability
Log[AM-101], µM
(C)
Diazepam
AM-101
% Viability
Log[Diazepam], µM
(D)
Control
Diazepam (20 µM)
AM-101 (3 µM)
10x
10x
10x

Figure S3

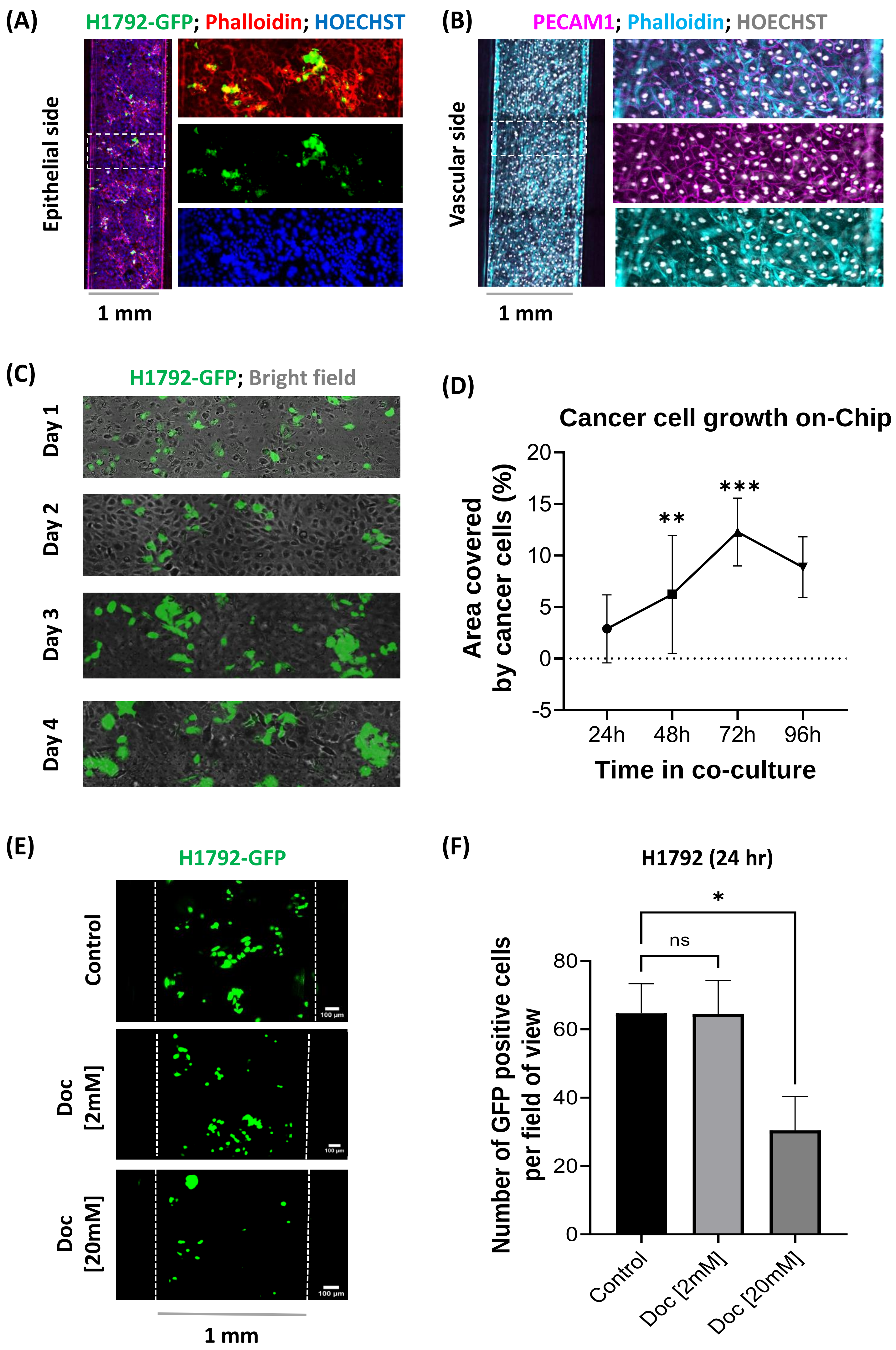
(A)
H1792-GFP; Phalloidin; HOECHST
Epithelial side
1 mm
(B)
PECAM1; Phalloidin; HOECHST
Vascular side
1 mm
(C)
H1792-GFP; Bright field
Day 1
Day 2
Day 3
Day 4
(D)
Cancer cell growth on-Chip
Area covered by cancer cells (%)
20
15
10
5
0
-5
**
***
24h
48h
72h
96h
Time in co-culture
(E)
H1792-GFP
Control
Doc [2mM]
Doc [20mM]
100 µm
1 mm
(F)
H1792 (24 hr)
Number of GFP positive cells per field of view
80
60
40
20
0
ns
*
Control
Doc [2mM]
Doc [20mM]

### Figure S4

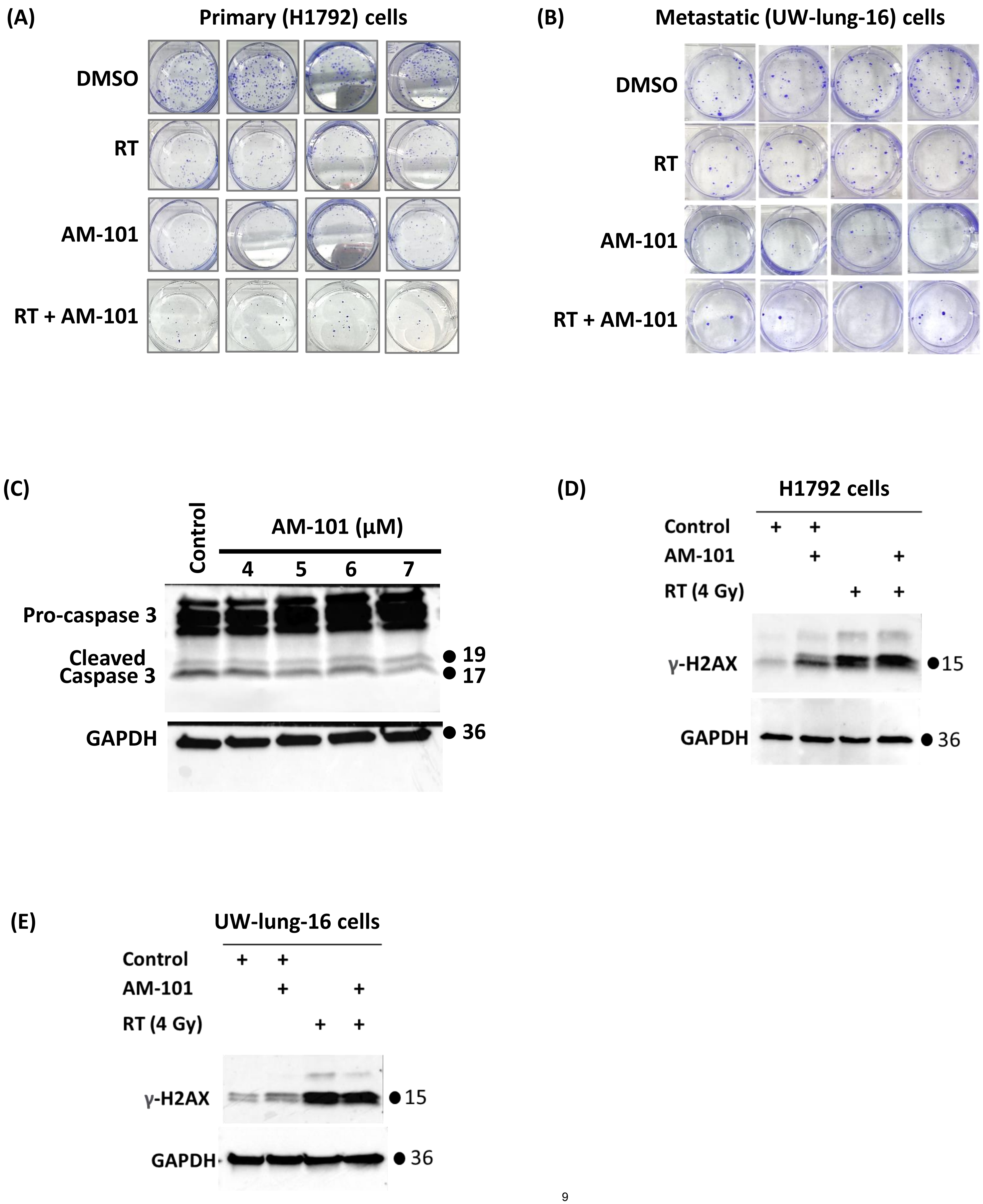
(A)
Primary (H1792) cells
DMSO
RT
AM-101
RT + AM-101
(B)
Metastatic (UW-lung-16) cells
DMSO
RT
AM-101
RT + AM-101
(C)
Control
AM-101 (µM)
4
5
6
7
Pro-caspase 3
Cleaved Caspase 3
19
17
GAPDH
36
(D)
H1792 cells
Control
AM-101
RT (4 Gy)
γ-H2AX
15
GAPDH
36
(E)
UW-lung-16 cells
Control
AM-101
RT (4 Gy)
γ-H2AX
15
GAPDH
36

### Figure S5

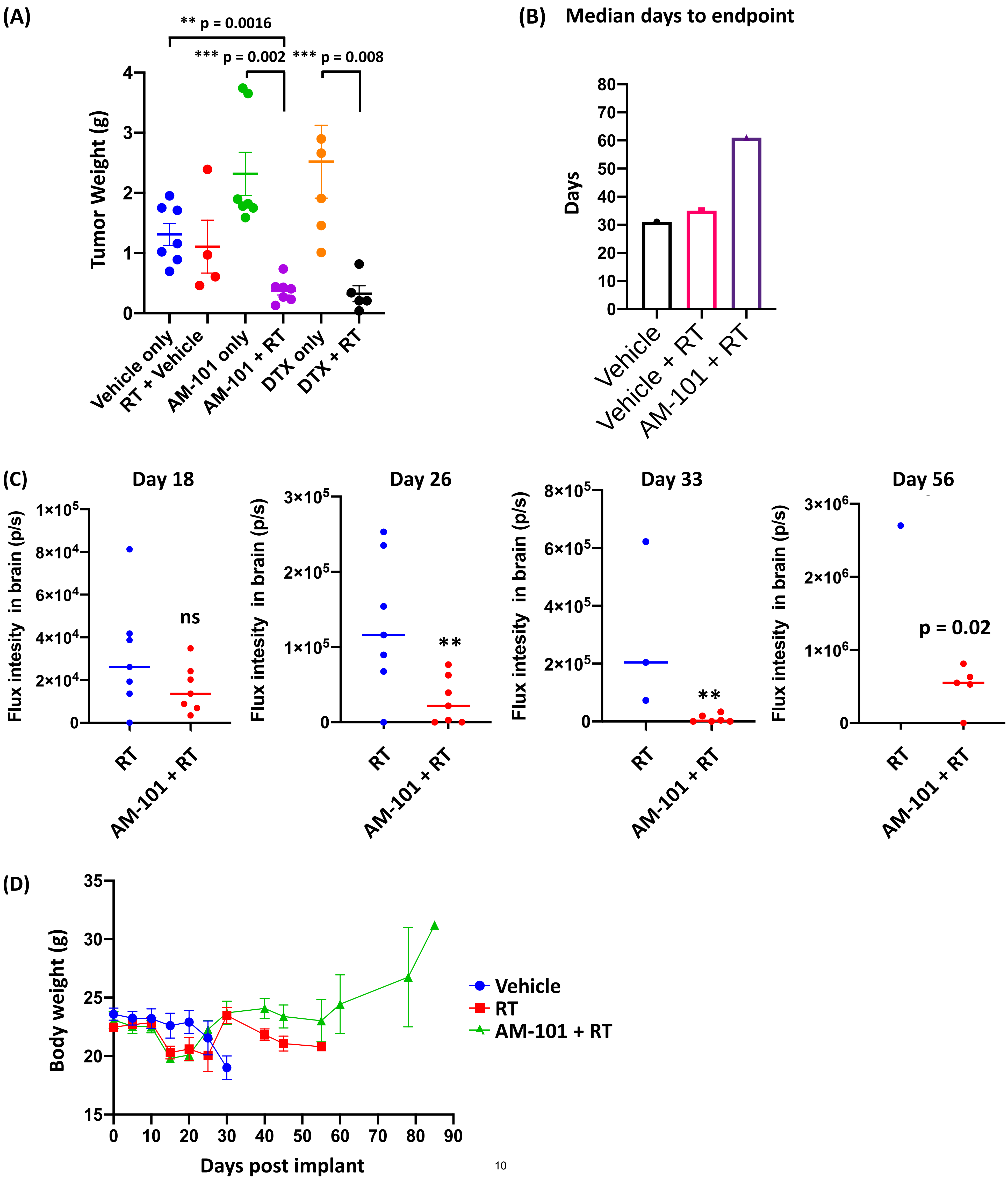

(A)
** p = 0.0016
*** p = 0.002
*** p = 0.008
Tumor Weight (g)
Vehicle only
RT + Vehicle
AM-101 only
AM-101 + RT
DTX only
DTX + RT
(B) Median days to endpoint
Days
Vehicle
Vehicle + RT
AM-101 + RT
(C)
Day 18
Day 26
Day 33
Day 56
Flux intesity in brain (p/s)
ns
**
p = 0.02
RT
AM-101 + RT
(D)
Body weight (g)
Days post implant
Vehicle
RT
AM-101 + RT

Figure S6

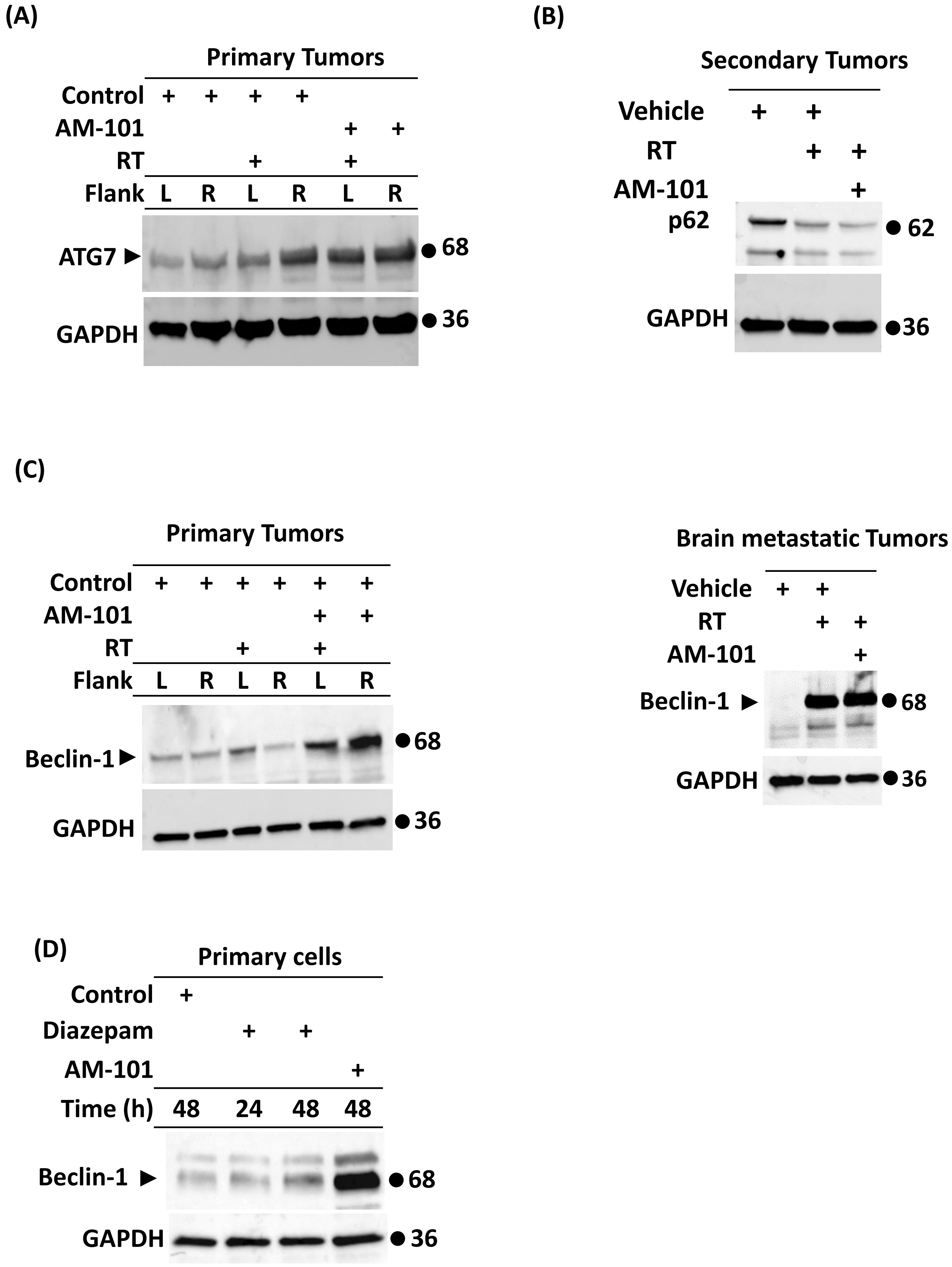
(A)
Primary Tumors
Control + + + +
AM-101 + +
RT + +
Flank L R L R L R
ATG7
68
GAPDH
36
(B)
Secondary Tumors
Vehicle + +
RT + +
AM-101 +
p62
62
GAPDH
36
(C)
Primary Tumors
Control + + + + + +
AM-101 + +
RT + +
Flank L R L R L R
Beclin-1
68
GAPDH
36
Brain metastatic Tumors
Vehicle + +
RT + +
AM-101 +
Beclin-1
68
GAPDH
36
(D)
Primary cells
Control +
Diazepam + +
AM-101 +
Time (h) 48 24 48 48
Beclin-1
68
GAPDH
36

**Figure S7**

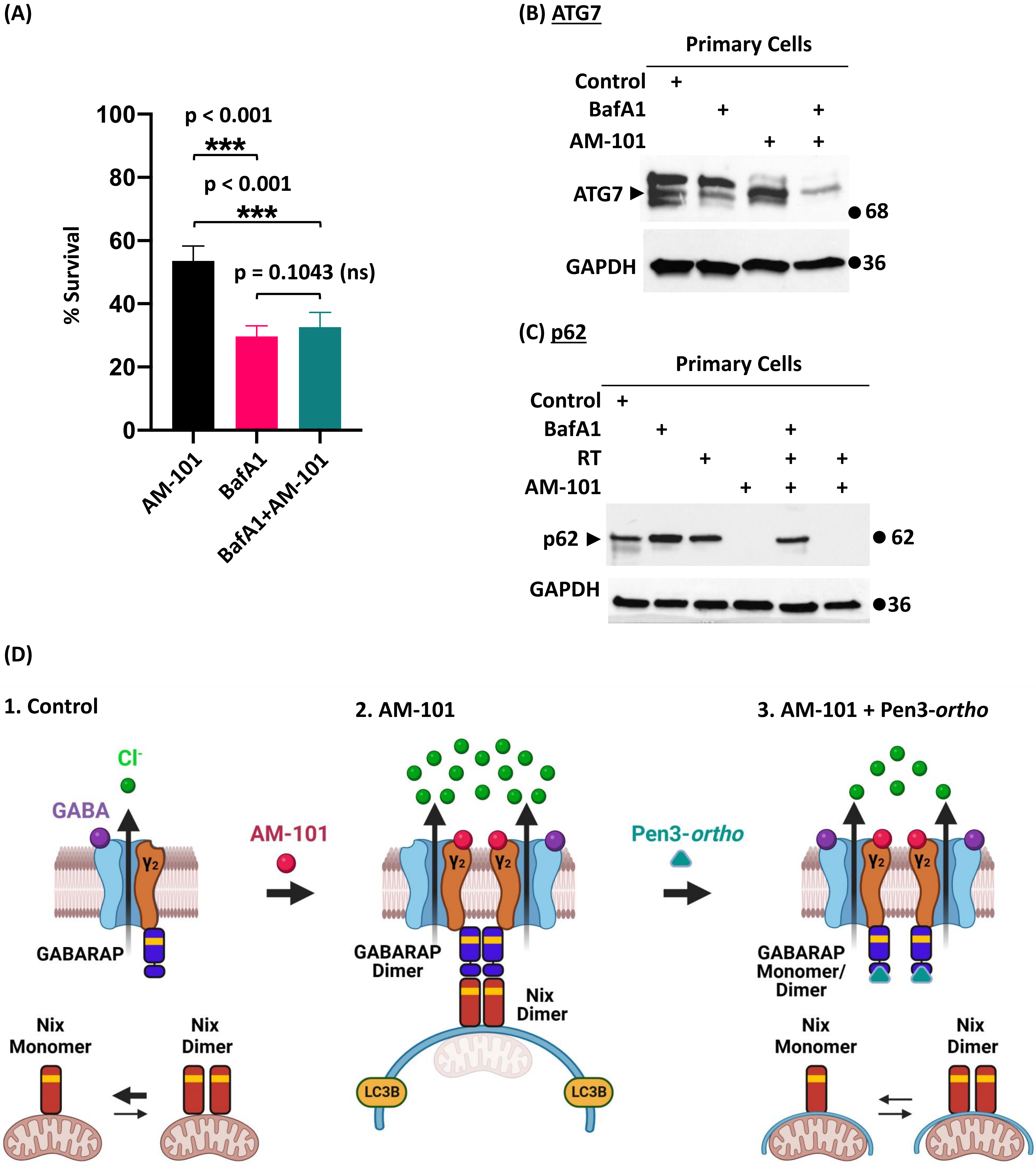
(A)
100
80
60
40
20
0
% Survival
p < 0.001
***
p < 0.001
***
p = 0.1043 (ns)
AM-101
BafA1
BafA1+AM-101
(B) ATG7
Primary Cells
Control
BafA1
AM-101
ATG7
68
GAPDH
36
(C) p62
Primary Cells
Control
BafA1
RT
AM-101
p62
62
GAPDH
36
(D)
1. Control
Cl-
GABA
γ2
GABARAP
Nix Monomer
Nix Dimer
AM-101
2. AM-101
γ2
γ2
GABARAP Dimer
Nix Dimer
LC3B
LC3B
Pen3-ortho
3. AM-101 + Pen3-ortho
γ2
γ2
GABARAP Monomer/ Dimer
Nix Monomer
Nix Dimer
